# Melanocortin 1 receptor regulates cholesterol and bile acid metabolism in the liver

**DOI:** 10.1101/2022.11.08.515543

**Authors:** Keshav Thapa, James J. Kadiri, Karla Saukkonen, Iida Pennanen, Minying Cai, Eriika Savontaus, Petteri Rinne

**Author notes:** Corresponding author: Petteri Rinne, PhD, Institute of Biomedicine, University of Turku, Kiinamyllynkatu 10, 20520 Turku, Finland.

## Abstract

Melanocortin 1 receptor (MC1-R) is widely expressed in melanocytes and leukocytes, and is thus strongly implicated in the regulation of skin pigmentation and inflammation. MC1-R mRNA has also been found in the rat and human liver, but its functional role has remained elusive. We hypothesized that MC1-R is functionally active in the liver and involved in the regulation of cholesterol and bile acid metabolism. We generated hepatocyte-specific MC1-R knock-out (*L-Mc1r*^*-/-*^) mice and phenotyped the mouse model for lipid profiles, liver histology and bile acid levels. *L-Mc1r*^*-/-*^ mice had significantly increased liver weight, which was accompanied by elevated levels of total cholesterol and triglycerides in the liver as well as in the plasma. These mice demonstrated also enhanced liver fibrosis and a disturbance in bile acid metabolism as evidenced by markedly reduced bile acid levels in the plasma and feces. Mechanistically, using HepG2 cells as an *in vitro* model, we found that selective activation of MC1-R in HepG2 cells reduced cellular cholesterol content and enhanced uptake of low- and high-density lipoprotein particles *via* a cAMP-independent mechanism. In conclusion, the present results demonstrate that MC1-R signaling in hepatocytes regulates cholesterol and bile acid metabolism and its deficiency leads to hypercholesterolemia and enhanced lipid accumulation and fibrosis in the liver.

## Introduction

Obesity is recognized as a global epidemic and is a major risk factor for type 2 diabetes, dyslipidemia and cardiovascular disease (B. Caballero, 2007). In particular, visceral obesity is associated with atherogenic dyslipidemia characterized by high levels of triglycerides (TG), TG-rich lipoproteins and low-density lipoprotein (LDL) cholesterol and reduced of high-density lipoprotein (HDL) cholesterol in the blood (Grundy, 2004; Klop et al., 2013). The liver plays a central role in this pathogenetic process as a regulator of cholesterol and fatty acid metabolism. As a consequence of hepatic dysregulation, *de novo* production and storage of cholesterol and fatty acids are enhanced leading to lipid accumulation in the liver, which eventually manifests as non-alcoholic fatty liver disease (NAFLD) (Diehl & Day, 2017; Friedman et al., 2018; Than & Newsome, 2015). Novel therapeutic strategies to enhance clearance and reduce excessive production of cholesterol and fatty acids in the liver are needed to mitigate the burden of obesity-associated dyslipidemia, NAFLD and associated cardiovascular complications such as atherosclerosis.

Melanocortins are a family of peptide hormones that are proteolytically cleaved from the precursor molecule proopiomelanocortin (POMC) to yield adrenocorticotrophin (ACTH) and α-, β-, and γ-melanocyte stimulating hormone (α-, β- and γ-MSH) (Gantz & Fong, 2003; Smith & Funder, 1988). Melanocortins bind to and activate five different G-protein coupled melanocortin receptor subtypes named from MC1-R to MC5-R (Gantz & Fong, 2003). Melanocortins and their receptors are expressed in the brain as well as in the periphery and regulate important physiological functions including skin pigmentation, sexual behavior, immune responses and energy homeostasis (Catania et al., 1996; Wikberg & Mutulis, 2008; Yeo et al., 2021). There has been wide interest in melanocortin receptors as potential drug targets and in fact, melanocortin receptor-targeted drugs have been recently approved for the treatment of rare skin diseases and genetic obesity syndromes (Montero-Melendez et al., 2022). MC1-R was the first member of the melanocortin receptor family to be cloned and it binds only α-MSH with high affinity (Mountjoy, 1994; Mountjoy et al., 1992). MC1-R is abundantly expressed in melanocytes in the skin and thus implicated as an integral regulator of skin pigmentation. MC1-R expression has been also demonstrated on a variety of peripheral cells including monocytes, macrophages, dendritic cells, neutrophils, endothelial cells and fibroblasts (Becher et al., 1999; Catania et al., 1996; Hartmeyer et al., 1997; Reichrath et al., 2005). Accordingly, increasing evidence demonstrates that MC1-R mediates potent and wide-ranging anti-inflammatory actions by suppressing the production of pro-inflammatory cytokines while simultaneously increasing the production of anti-inflammatory cytokines (Catania et al., 2004). Intriguingly, MC1-R mRNA has also been detected in the human and rat liver following the testing of a hypothesis that MC-Rs might modulate inflammation in the liver (Gatti et al., 2006; Malik et al., 2012). However, these early studies aimed to only characterize the expression profile of different MC-Rs in the liver and the functional role of hepatic MC1-R has thereby remained unexplored.

We have recently found that global deficiency of MC1-R signaling accelerates atherosclerosis in apolipoprotein E knockout mice (Apoe^-/-^) by increasing arterial monocyte accumulation and by disturbing cholesterol and bile acid metabolism (Rinne et al., 2018). Specifically, MC1-R deficient mice showed elevated cholesterol levels in the plasma and liver in conjunction with a distinct bile acid profile characterized by reduced primary and increased secondary bile acid levels. This phenotype could be however caused by multiple mechanisms leaving an open question whether MC1-R in the liver has a regulatory role in cholesterol and bile acid metabolism. In the present study, we aimed to address this question by engineering a hepatocyte-specific MC1-R knock-out mouse model. We here show that the loss of MC1-R signaling in hepatocytes causes hypercholesterolemia and enhanced lipid accumulation in the liver, and disturbs bile acid metabolism. Moreover, using HepG2 cells as an *in vitro* model, we found that α-MSH and selective MC1-R activation reduced cellular cholesterol content and enhanced uptake of LDL and HDL. This study demonstrates that MC1-R is functionally active in the liver and regulates cholesterol and bile acid metabolism in a protective way that could also have therapeutic implications.

## Results

### Hepatocyte-specific MC1-R deficiency enhances cholesterol and lipid accumulation in the liver

We first aimed to investigate whether MC1-R is expressed in the mouse liver. Immunohistochemical staining revealed a strong and uniform expression of MC1-R in hepatocytes (**Figure 1A**). Furthermore, we sought to investigate whether the expression level of MC1-R in the liver is affected by feeding mice a cholesterol-rich Western diet. Remarkably, 12 weeks of Western diet feeding resulted in significant downregulation of MC1-R mRNA level in the liver (**Figure 1B**). This result was further corroborated by Western blotting, which showed reduced protein expression of MC1-R in the liver of Western diet-fed mice (**Figure 1C**). The specificity of the MC1-R signal was validated by pre-adsorption of the antibody with a MC1-R blocking peptide (**Supplementary Figure 1**).

**Figure 1.**
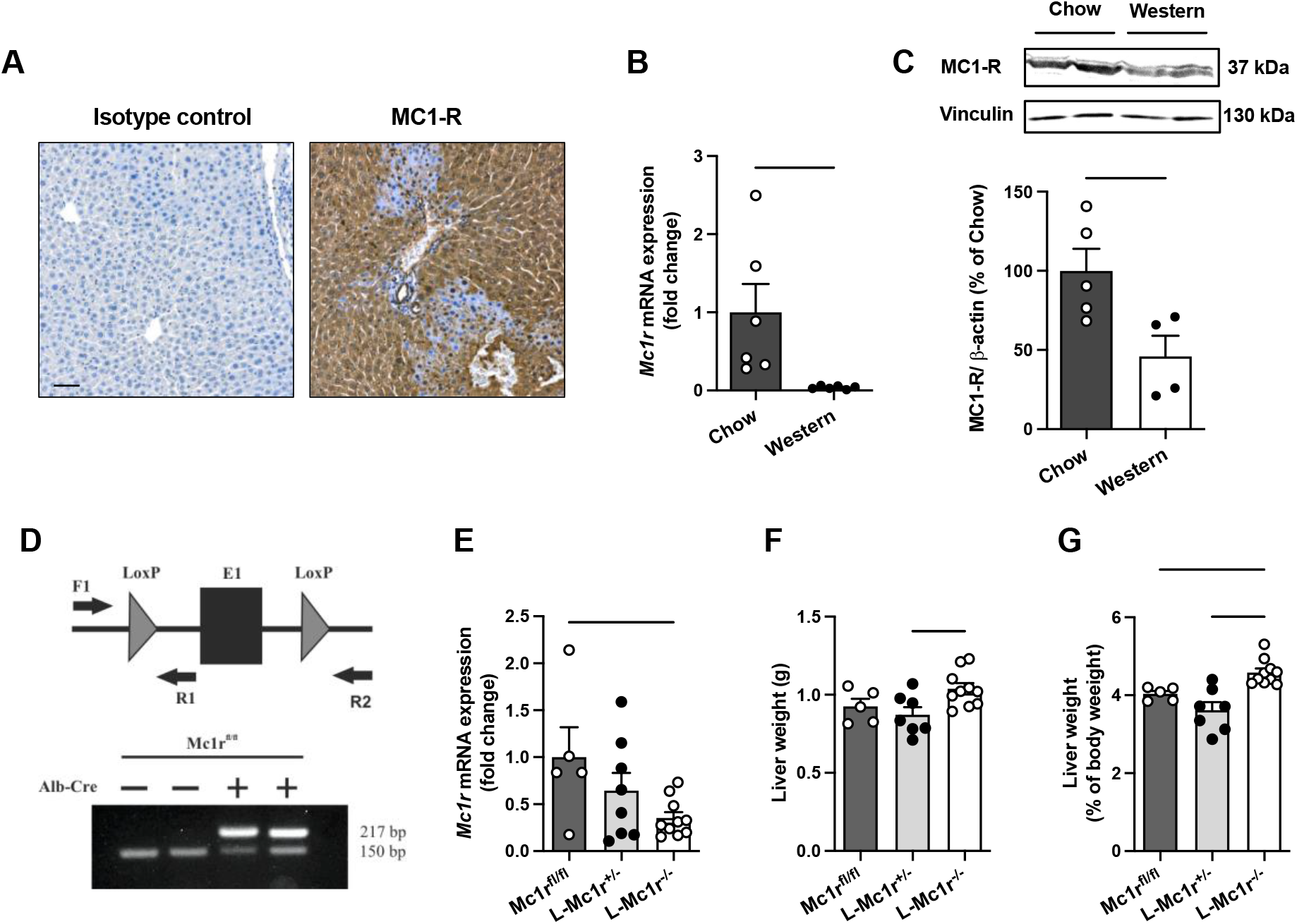
MC1-R is expressed in the mouse liver and down-regulated in mice fed a cholesterol-rich diet. (**A)** Immunostaining of MC1-R staining in the liver of chow-fed C57Bl/6J mouse. In control section, anti MC1-R antibody was replaced by purified normal rabbit IgG (isotype control). Scale bar, 50 μm. (**B)** Quantitative real-time polymerase chain reaction (qPCR) analysis of *Mc1r* mRNA expression in the liver of chow- and Western diet-fed mice. (**C)** Representative Western blots of MC1-R and β-actin (loading control) and quantification of MC1-R protein level in the liver of chow- and Western diet-fed mice. (**D)** Schematic presentation of the loxP-flanked (floxed) *Mc1r* allele and the positions of forward and reverse primers used for PCR genotyping. PCR analysis of genomic DNA extracted from the liver of Alb-Cre-negative and -positive mice that were homozygous for the Mc1r floxed allele (Mc1r^fl/fl^). The size of the recombined allele is ∼217 bp. (**E**) qPCR analysis of *Mc1r* expression in the liver of chow-fed *Mc1r*^*fl/fl*^, *L-Mc1r*^*+/-*^ and *L-Mc1r*^*-/-*^ mice at the age of 16 weeks. (**F, G**) Absolute liver weight and liver to body weight ratio (expressed as percentage of body weight) in chow-fed *Mc1r*^*fl/fl*^, *L-Mc1r*^*+/-*^ and *L-Mc1r*^*-/-*^ mice at the age of 16 weeks. Values are mean ± SEM, n = 5-10 mice per group in each graph. \**P<0*.*05* and \*\**P<0*.*01* for the indicated comparisons by one-way ANOVA and Dunnet *post hoc* tests. *L-Mc1r*^*+/-*^ indicates Alb-Cre^+/-^ mice that were heterozygous for the *Mc1r* floxed allele; *L-Mc1r*^*-/-*^, hepatocyte-specific MC1-R knock-out mice.

To determine the regulatory role of MC1-R in the liver, we generated hepatocyte-specific MC1-R knock-out mice (denoted as *L-Mc1r*^*-/-*^) by crossing MC1-R floxed (*Mc1r*^*fl/fl*^) mice with transgenic mice expressing Cre recombinase under the control of the mouse albumin promoter (Alb-Cre^+/-^) (**Figure 1D**). Genotyping of the liver samples verified efficient recombination of the loxP-flanked allele in *L-Mc1r*^*-/-*^ mice (**Figure 1E**) that resulted in significant downregulation of hepatic *Mc1r* mRNA expression in these mice compared to control Mc1r^fl/fl^ mice (**Figure 1E**). A gene dosage effect was also noted in this regard as Alb-Cre^+/-^ mice that were heterozygous for the *Mc1r* floxed allele (denoted as *L-Mc1r*^*+/-*^) showed only partial downregulation of *Mc1r* compared to *L-Mc1r*^*-/-*^ mice (**Figure 1E**). To evaluate the effect of hepatocyte-specific MC1-R deficiency on body weight development, *Mc1r*^*fl/fl*^, *L-Mc1r*^*+/-*^ and *L-Mc1r*^*-/-*^ mice were fed a normal chow diet and weighed weekly during a monitoring period from 8 to 16 weeks of age. However, no differences were observed in body weight between the genotypes (**Supplementary Figure 2A**). Body composition analysis by quantitative NMR scanning at the start and end of the monitoring period did not reveal any significant changes in total fat or lean mass of *L-Mc1r*^*-/-*^ mice compared to control genotypes (**Supplementary Figure 2B, C**). Of note, *L-Mc1r*^*-/-*^ mice displayed a significant increase in liver weight (**Figure 1F**), which became more evident when calculated as percentage of body weight (**Figure 1G**). Since the relative liver weight was significantly increased in comparison with both control groups, we used *L-Mc1r*^*+/-*^ mice as the control group in subsequent analyses to eliminate the possible confounding by Alb-Cre transgene expression.

Histological examination by H&E and Oil Red O staining revealed enhanced accumulation of intracellular lipid droplets in the liver of *L-Mc1r*^*-/-*^ mice in comparison to the control *L-Mc1r*^*+/-*^ mice **(Figure 2A)**. Supporting this finding, quantification of hepatic lipid content showed increased triglyceride (TG) and total cholesterol levels in *L-Mc1r*^*-/-*^ mice **(Figure 2B, C)**. Likewise, plasma TG and TC levels were significantly higher in *L-Mc1r*^*-/-*^ mice compared to control *L-Mc1r*^*+/-*^ mice **(Figure 2D, E**). *L-Mc1r*^*-/-*^ mice also demonstrated signs of increased liver fibrosis, as evidenced by Picrosirius Red staining and gene expression analysis of fibrotic genes (**Supplementary Figure 3A-C)**. However, no change in the expression of pro-inflammatory genes was observed between the genotypes (**Supplementary Figure 3D)**.

**Figure 2.**
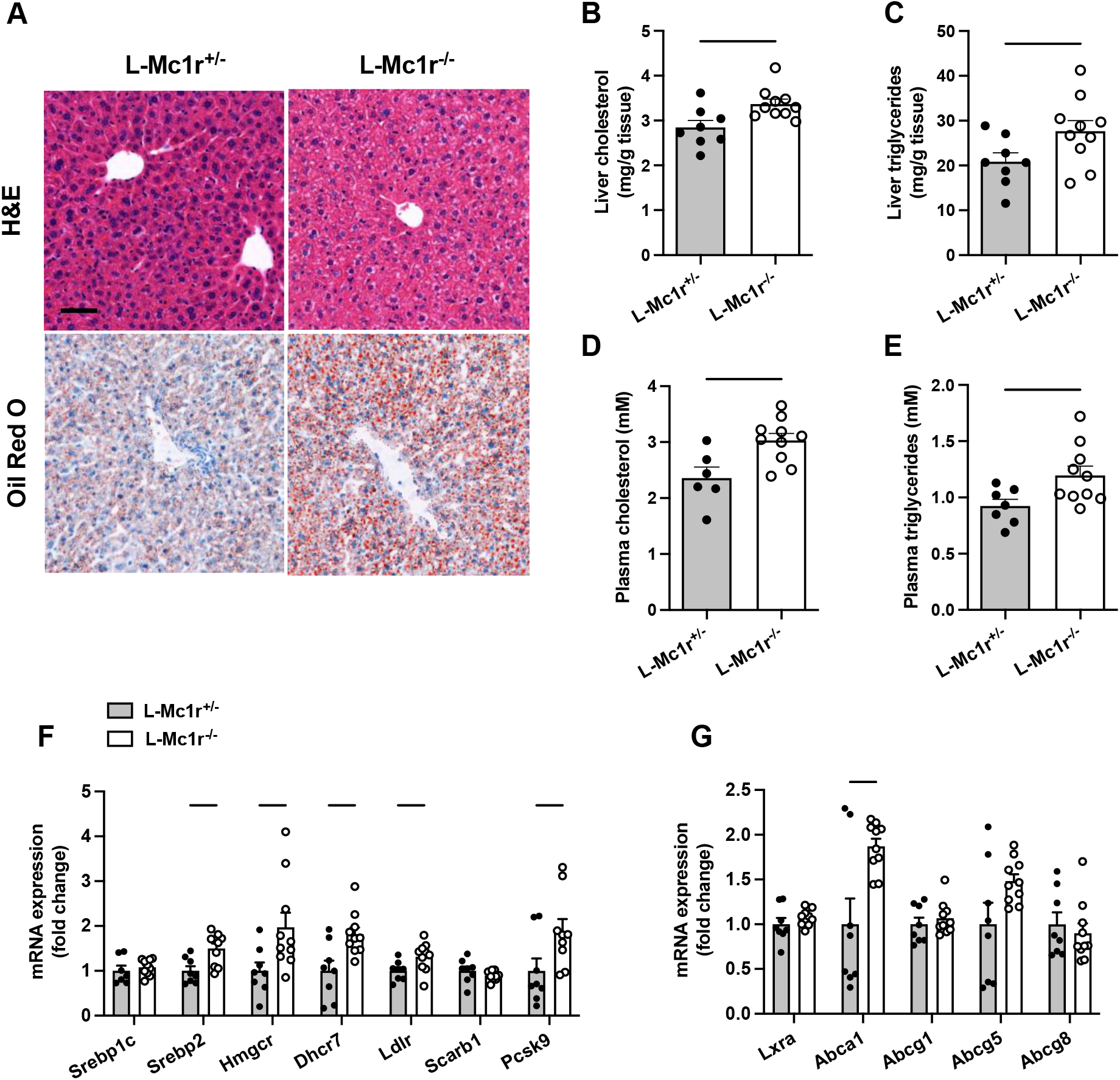
Hepatocyte-specific MC1-R deficiency enhances cholesterol and triglyceride accumulation in the liver. (**A**) Representative hematoxylin and eosin (H&E) and Oil Red O-stained liver sections of chow-fed *L-Mc1r*^*+/-*^ and *L-Mc1r*^*-/-*^ mice. Scale bar, 50 μm. (**B, C**) Quantification of liver total cholesterol and triglyceride content in chow-fed *L-Mc1r*^*+/-*^ and *L-Mc1r*^*-/-*^ mice. (**D, E**) Quantification of plasma total cholesterol and triglyceride concentrations in chow-fed *L-Mc1r*^*+/-*^ and *L-Mc1r*^*-/-*^ mice. **(F, G)** qPCR analysis of cholesterol synthesis and transporter genes and their transcriptional regulators in the liver of chow-fed *L-Mc1r*^*+/-*^ and *L-Mc1r*^*-/-*^ mice. Values are mean ± SEM, n = 6-10 mice per group in each graph. \**P<0*.*05* and \*\**P<0*.*01* versus *Mc1r*^*+/-*^ mice by Student’s t test.

To identify molecular mechanisms behind the increased hepatic lipid accumulation and hypercholesterolemia, we quantified the expression levels of genes involved in cholesterol synthesis and transport. We found that sterol regulatory element binding protein 2 (*Srebp2*), which is the master transcriptional regulator of cholesterol homeostasis, was upregulated in the liver of *L-Mc1r*^*-/-*^ mice, while the expression of *Srebp1c* that primarily regulates fatty acid synthesis was unchanged **(Figure 2F)**. Consequently, SREBP2 target genes *Hmgcr* and *Dhcr7*, which are crucially involved in the biosynthesis of cholesterol, were significantly upregulated in *L-Mc1r*^*-/-*^ mice **(Figure 2F)**. We also looked at the mRNA expression levels of lipoprotein receptors and found that LDL receptor (*Ldlr*) mRNA levels were increased in *L-Mc1r*^*-/-*^ mice, but no change was observed on the HDL receptor SR-BI (*Scarb1*) expression (**Figure 2F**). The change in *Ldlr* expression was paralleled by a tendency (P=0.07) towards increased expression of proprotein convertase subtilisin/kexin type 9 (*Pcsk9*) that promotes lysosomal degradation of LDL receptor in the liver and is also transcriptionally regulated by SREBP2 **(Figure 2F)**. Aside from genes involved in cholesterol synthesis and uptake, there was no genotype difference in the mRNA level of liver X receptor alpha (*Lxra*) that induces transcription of genes that control cellular cholesterol efflux **(Figure 2G)**. However, among different cholesterol efflux transporters, the ATP-binding cassette transporter *Abca1* was significantly upregulated and *Abcg5* expression tended to be increased (P=0.05) in the *L-Mc1r*^*-/-*^ mice, while no change was observed in *Abcg1* or *Abcg8* expression level **(Figure 2G)**.

### Hepatocyte-specific MC1-R deficiency disturbs bile acid metabolism

Based on the previous finding of disturbed bile acid metabolism in global MC1-R deficient mice on Apoe^-/-^ background, we were curious to investigate whether the hepatocyte-specific MC1-R knockout model recapitulates this phenotype. To this end, we quantified total and individual bile acids (BA) in the liver, feces and plasma of *L-Mc1r*^*-/-*^ mice by liquid chromatography-mass spectrometry. We found that the total amount of BAs was markedly reduced in the plasma and to some extent (P=0.06) also in the feces of *L-Mc1r*^*-/-*^ mice **(Figure 3B, C)**, while the size of hepatic BA pool remained unchanged **(Figure 3A)**. These changes were largely attributable to reduction in secondary BAs **(Figure 3B, C)**. Quantification of primary BA species in the plasma revealed that the levels of taurine-conjugated cholic acid (CA) and ursodeoxycholic acid (UDCA) were lower in *L-Mc1r*^*-/-*^ mice. In terms of secondary BAs, *L-Mc1r*^*-/-*^ mice showed significantly reduced plasma levels of taurine-conjugated deoxycholic acid (DCA), hyodeoxycholic acid (HDCA) and ω-muricholic acid (ω-MCA) **(Figure 3E)**. The amount of DCA was also lower in the liver of *L-Mc1r*^*-/-*^ mice, while in the feces, HDCA, litocholic acid (LCA) and 12-keto litocholic acid (12-oxo LCA), which is the primary metabolite of DCA, were significantly reduced by MC1-R deficiency (**Supplementary Figure 4**). Furthermore, the relative proportions of primary BAs in the plasma indicate that hepatocyte-specific MC1-R deficiency reduced the amount of CA with an accompanying increase in CDCA and UDCA (**Figure 3F**). This BA profile is further reflected as a significant reduction in the plasma ratio of CA:CDCA **(Figure 3G)**.

To address possible causes of disturbed BA metabolism in *L-Mc1r*^*-/-*^ mice, we quantified the hepatic expression of genes encoding for BA synthetizing enzymes **(Figure 4A)**. Although the expression of cholesterol 7 alpha-hydroxylase (encoded by *Cyp7a1*), which is the first and rate-liming enzyme in BA synthesis, was unchanged, *L-Mc1r*^*-/-*^ mice demonstrated significant upregulation of sterol 12α-hydroxylase (encoded by *Cyp8b1*) and sterol 27-hydroxylase (encoded by *Cyp27a1*) **(Figure 4A)**. Furthermore, *L-Mc1r*^*-/-*^ mice had reduced mRNA level of steroidogenic acute regulatory protein 1 (*Stard1*), which facilitates trafficking of cholesterol to mitochondria and thus feeds the alternative mitochondrial pathway of BA synthesis **(Figure 4A)**. Secondly, we quantified the hepatic mRNA levels of transporters responsible for the uptake of BAs and their excretion into bile and systemic circulation **(Figure 4B)**. We found that the expression of sodium/bile acid cotransporter (encoded by *Ntcp*), which accounts for the majority (∼90%) of BA uptake from the portal circulation, was upregulated in the liver of *L-Mc1r*^*-/-*^ mice, while bile salt export pump (*Bsep*) showed no change at the mRNA level **(Figure 4B)**. An alternative basolateral export of BAs is mediated by the heterodimeric organic solute transporter OSTα/OSTβ and the multidrug resistance-associated proteins MRP3 and MRP4, the last of which was downregulated in *L-Mc1r*^*-/-*^ mice **(Figure 4B)**. Thirdly, among different nuclear receptors that regulate the transcription of BA enzymes and transporters, farnesoid X receptor (*Fxr*) and hepatocyte nuclear factor 4α (*Hnf4a*) were significantly upregulated in the liver of *L-Mc1r*^*-/-*^ mice **(Figure 4C)**.

**Figure 3.**
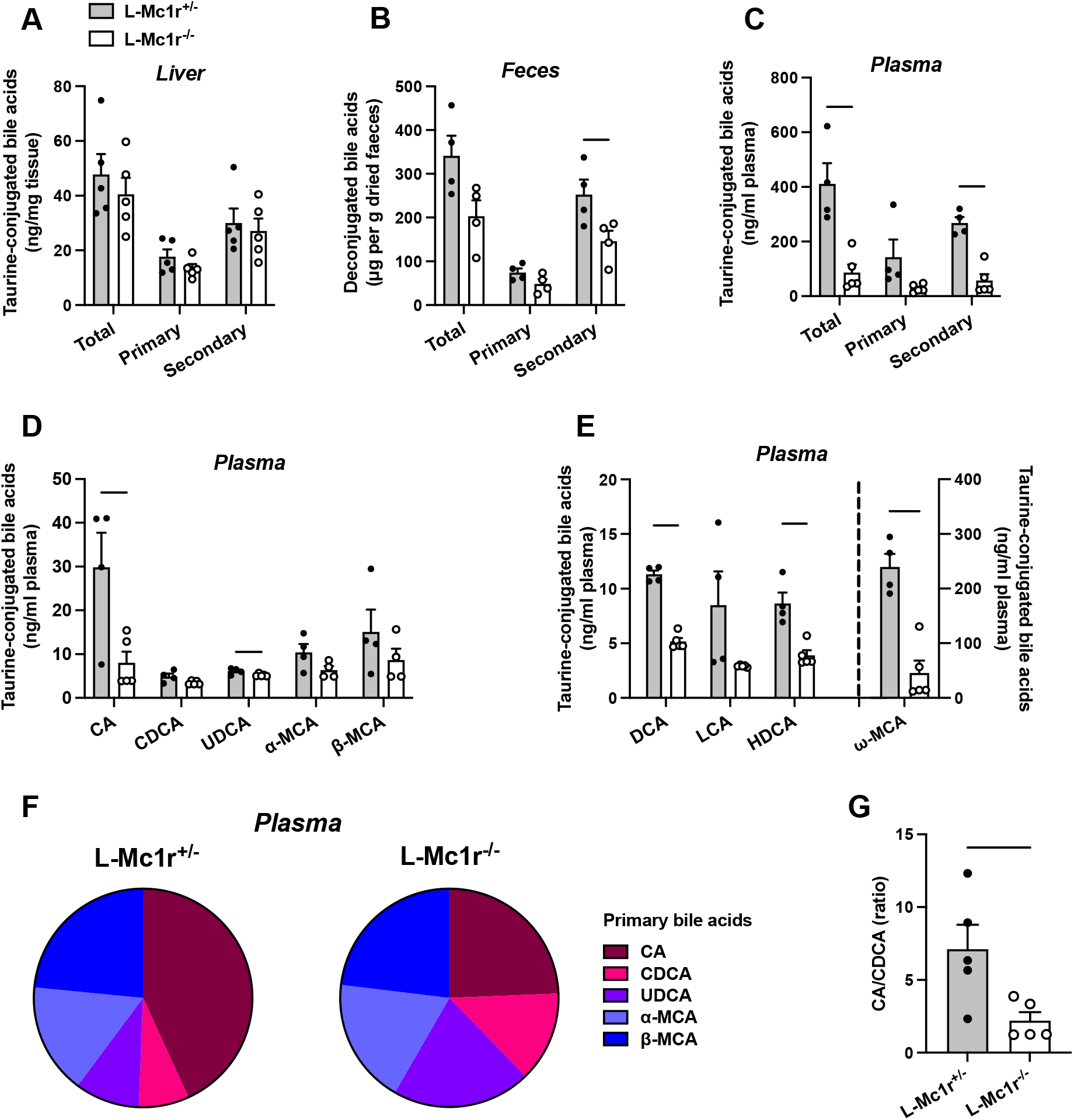
Hepatocyte-specific MC1-R deficiency disturbs bile acid metabolism. (**A-C**) Quantification of total, primary and secondary bile acids in the liver, plasma and feces of chow-fed *L-Mc1r*^*+/-*^ and *L-Mc1r*^*-/-*^ mice. (**D**) Quantification of individual primary bile acids in the plasma of chow-fed *L-Mc1r*^*+/-*^ and *L-Mc1r*^*-/-*^ mice. (**E**) Quantification of individual secondary bile acids in the plasma of chow-fed *L-Mc1r*^*+/-*^ and *L-Mc1r*^*-/-*^ mice. **(F)** Relative proportions of individual primary bile acids in the plasma of chow-fed *L-Mc1r*^*+/-*^ and *L-Mc1r*^*-/-*^ mice. (**G**) The ratio of cholic acid (CA) to chenodeoxycholic acid (CDCA) in the plasma of chow-fed *L-Mc1r*^*+/-*^ and *L-Mc1r*^*-/-*^ mice. Values are mean ± SEM, n = 4-5 mice per group in each graph. \**P<0*.*05*, \*\**P<0*.*01*, \*\*\**P<0*.*001* and \*\*\*\**P<0*.*0001* versus *L-Mc1r*^*+/-*^ mice by Student’s t test. CA indicates cholic acid; CDCA, chenodeoxycholic acid; UDCA, ursodeoxycholic acid; MCA, muricholic acid; DCA, deoxycholic acid; LCA, litocholic acid; HDCA, hyodeoxycholic acid (HDCA).

**Figure 4.**
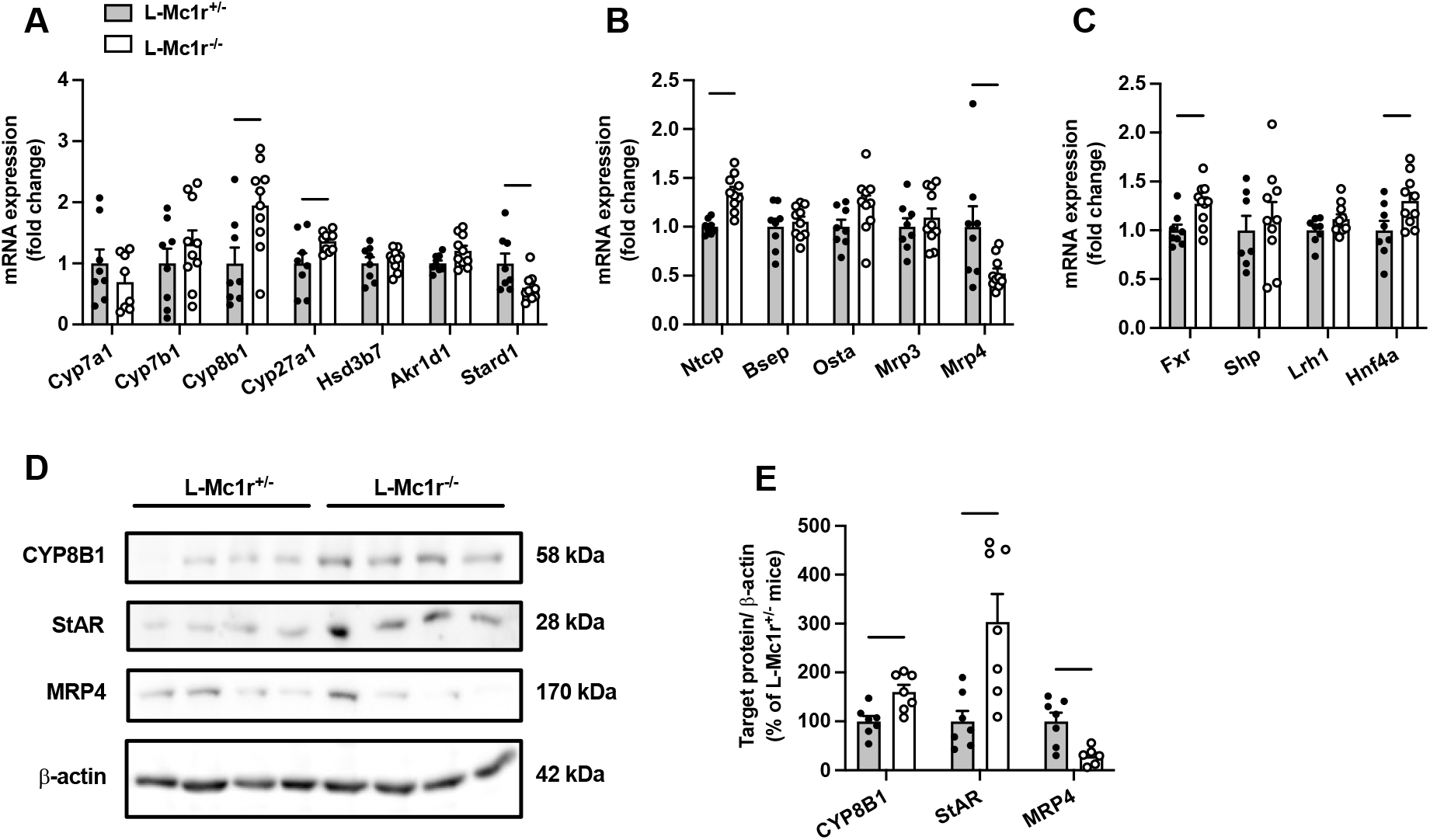
Hepatocyte-specific MC1-R deficiency affects the expression of genes involved in bile acid synthesis and transport. (**A, B**) qPCR analysis of genes involved in the bile acid synthesis and transport in the liver of chow-fed *Mc1r*^*fl/fl*^, *L-Mc1r*^*+/-*^ and *L-Mc1r*^*-/-*^ mice. (**C**) qPCR analysis of nuclear receptor genes that regulate the transcription of bile acid enzymes and transporters. (**D, E**) Representative Western blots and quantification of CYP8B1, StAR and MRP4 protein levels in the liver of chow- and Western diet-fed mice. Values are mean ± SEM, n = 7-10 mice per group in each graph. \**P<0*.*05*, \*\**P<0*.*01* and \*\*\**P<0*.*001* versus *Mc1r*^*+/-*^ mice by Student’s t test. *Cyp7a1*, cholesterol 7 alpha-hydroxylase; *Cyp7b1*, 25-hydroxycholesterol 7-alpha-hydroxylase; *Cyp8b1*, sterol 12-alpha-hydroxylase; *Cyp27a1*, sterol 27-hydroxylase; *Stard1*, steroidogenic acute regulatory protein; *Fxr*, farnesoid X receptor; *Lrh1*, liver receptor homologue 1; *Bsep*, bile-salt export pump; *Ntcp*, Na+-taurocholate cotransporting polypeptide; *Hnf4a*, hepatocyte nuclear factor 4 alpha.

Finally, we selected the genes, which were differently expressed in *L-Mc1r*^*-/-*^ mice and could potentially explain the observed BA profile in these mice, and quantified the corresponding protein levels of these gene products in the liver by Western blotting. In good agreement with the mRNA level changes, *L-Mc1r*^*-/-*^ mice showed higher CYP8B1 and lower MRP4 protein expression compared to control mice. However, StAR (encoded by *Stard1*) protein level was significantly increased in the liver of *L-Mc1r*^*-/-*^ mice, which contradicts the mRNA level finding.

### The endogenous MC1-R agonist α-MSH reduces cellular cholesterol content and enhances LDL and HDL uptake in HepG2 cells

The finding of MC1-R expression in the mouse liver and the phenotype of enhanced cholesterol accumulation in *L-Mc1r*^*-/-*^ mice led us to investigate the effects and underlying mechanisms of MC1-R activation in human HepG2 cells. First, we aimed to verify that human hepatocytes also express MC1-R. Indeed, HepG2 cells clearly express MC1-R protein (**Figure 5A**). Consistent with the finding of reduced MC1-R expression in the liver of Western diet-fed mice, loading of HepG2 cells with palmitic acid (a saturated free fatty acid) caused a rapid decrease of MC1-R protein expression (**Figure 5A**). However, exposure of HepG2 cells to excess LDL cholesterol (**Figure 5B**) or treatment with the HMGCR inhibitor atorvastatin (**Figure 5C**) to lower cellular cholesterol content did not change MC1-R protein level. Second, we studied how MC1-R activation affects cholesterol metabolism in HepG2 cells. For this purpose, HepG2 cells were stimulated with the endogenous MC1-R agonist α-MSH and the amount of cellular free cholesterol was quantified using Filipin staining. We observed that α-MSH (1 μM) significantly decreased the free cholesterol content in HepG2 cells with an effect appearing after 3 hours and plateauing towards the 24-hour time point (**Figure 5D**). In terms of concentration-responsiveness, cholesterol content was already reduced with a subnanomolar concentration (0.1 nM) of α-MSH and the maximal response was achieved with 1 μM α-MSH (**Figure 5E**). The reduction in cellular cholesterol was accompanied by significant increases in LDL and HDL uptake (**Figure 5F, G**), as evaluated after 24-hour treatment with α-MSH using fluorescently labeled lipoprotein particles (Dil-LDL and Dil-HDL). In good agreement with these findings, gene expression analysis revealed that α-MSH upregulated *LDLR* and *SCARB1* in a concentration-dependent manner (**Figure 5H, I**). These effects were only apparent at 3-hour time point, which probably reflects the short half-life of α-MSH. Nevertheless, upregulated *LDLR* and *SCARB1* mRNA levels translated into more sustained increases in the corresponding protein levels (LDLR and SR-BI) after α-MSH treatment (**Figure 5I, J**). Protein level analyses further showed that α-MSH had no effect on the expression of the cholesterol biosynthetic enzymes HMGCR and DHCR7 (**Figure 5J, K**). Intriguingly, we observed that LDLR expression at the cell surface, as quantified by flow cytometry, was markedly increased already after 1 hour treatment with 1 μM α-MSH (**Figure 5L**). Finally, in terms of BA metabolism, it appeared that α-MSH increased CA concentration in the culture medium of HepG2 cells without any effect on CDCA concentration (**Supplementary Figure 5A**). Consequently, the ratio of CA to CDCA was significantly increased in response to α-MSH treatment (**Supplementary Figure 5B**). Supporting this finding, Western blotting analysis showed that α-MSH upregulated CYP8B1 (**Supplementary Figure 5C**), which is the major determinant of CA:CDCA ratio.

**Figure 5.**
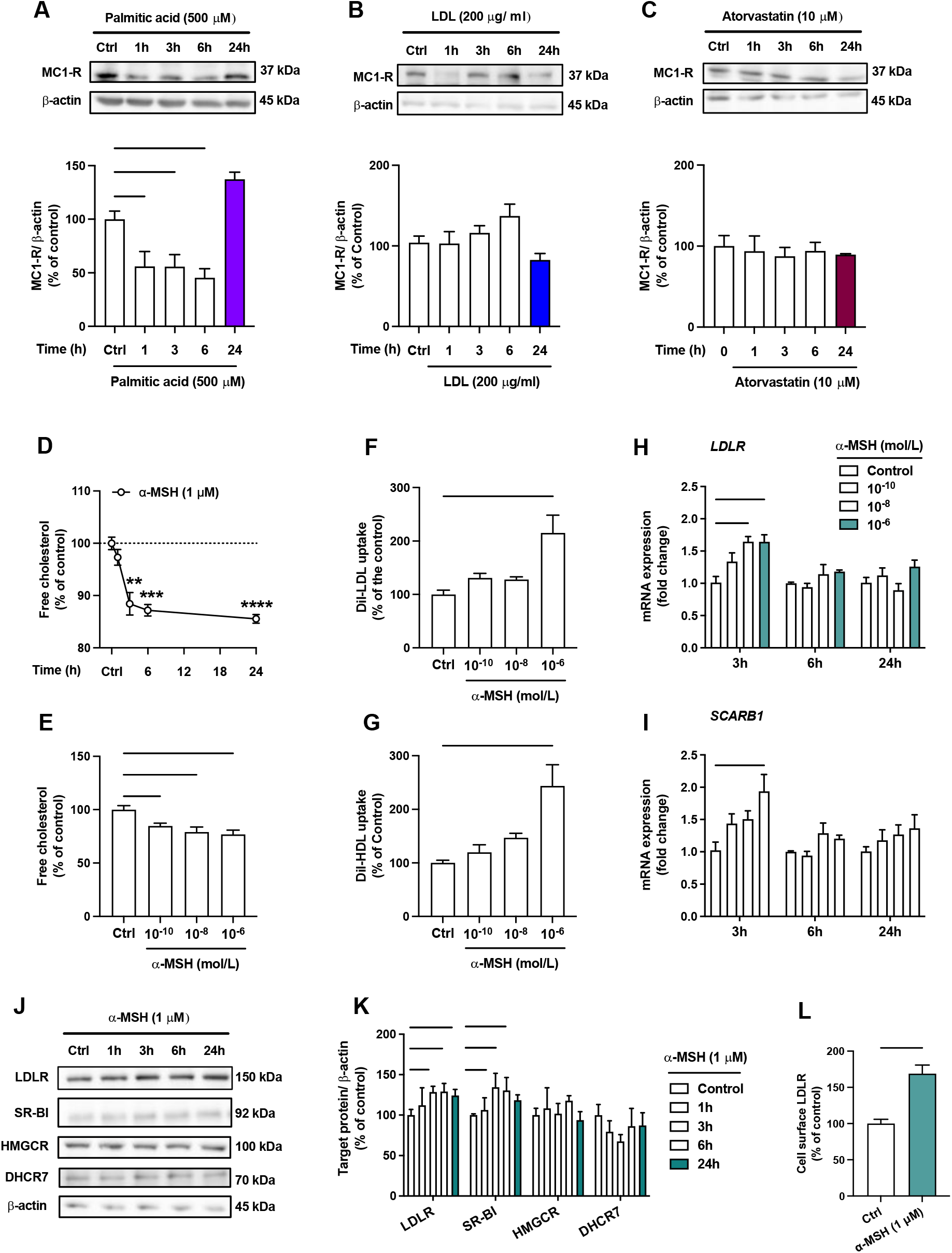
The endogenous MC1-R agonist α-MSH reduces cellular cholesterol content and enhances LDL and HDL uptake in HepG2 cells. (**A-C**) Representative Western blots and quantification of MC1-R protein level in HepG2 cells treated with palmitic acid (500 μM), LDL (200 μg/ml) or atorvastatin (10 μM) for 1, 3, 6 or 24 hours. **(D)** Quantification of free cholesterol content using filipin staining in HepG2 cells treated with α-MSH (1 μM) for 1, 3, 6 or 24 hours. **(E)** Quantification of free cholesterol content in HepG2 cells treated with different concentrations of α-MSH (0.1 nM, 10 nM or 1 μM) for 24 hours. **(F, G)** Quantification of LDL and HDL uptake in HepG2 cells treated with different concentrations (0.1 nM, 10 nM or 1 μM) of α-MSH for 24 hours. **(H, I)** qPCR analysis of *LDLR* and *SCARB1* expression in HepG2 cells treated with different concentrations of α-MSH for 3, 6 or 24 hours. **(J, K)** Representative Western blots and quantification of LDL-R and SR-BI proteins levels in HepG2 cells treated with 1μM α-MSH for 1, 3, 6 or 24 hours. (**L**) Quantification of cell surface LDLR by flow cytometry in HepG2 cells treated with 1μM α-MSH for 24 hours. Values are mean ± SEM, \**P<0*.*05* and \*\**P<0*.*01* for the indicated comparisons by one-way ANOVA and Dunnet *post hoc* tests (**A-K**) or by Student’s t test (**L**).

### Selective activation of MC1-R mimics the effects of α-MSH in HepG2 Cells

Since α-MSH can also bind and activate other MC-R subtypes, we aimed to verify that the effects induced by α-MSH were particularly derived from the activation of MC1-R. To this end, we repeated the key experiments using LD211, which is a highly potent and selective agonist for MC1-R with no detectable binding to other MC-R subtypes (Doedens et al., 2010). Closely mirroring the effect observed with α-MSH, selective activation of MC1-R with LD211 led to a concentration-dependent reduction in cellular cholesterol amount (**Figure 6A**). LD211 also significantly increased LDL and HDL uptake (**Figure 6B, C**), which was accompanied by upregulation of *LDLR* and *SCARB1* mRNA expression (**Figure 6D**). In comparison with α-MSH (**Figure 4G, H**), LD211 caused more sustained changes in gene expression (**Figure 6D**), which could be explained by the cyclic and more stable structure of LD211 (Doedens et al., 2010). Despite the clear-cut effects observed at the mRNA level, LD211 did not significantly affect LDLR and SR-BI protein levels, as detected by Western blotting. However, the cell surface expression of LDLR, which is a major determinant of LDL uptake rate, was markedly increased in HepG2 cells after 3-, 6- and 24-hour treatment with LD211 (**Figure 6E**). Taken together, these results demonstrate that the α-MSH-induced effects were largely reproducible by selective MC1-R activation with LD211.

**Figure 6.**
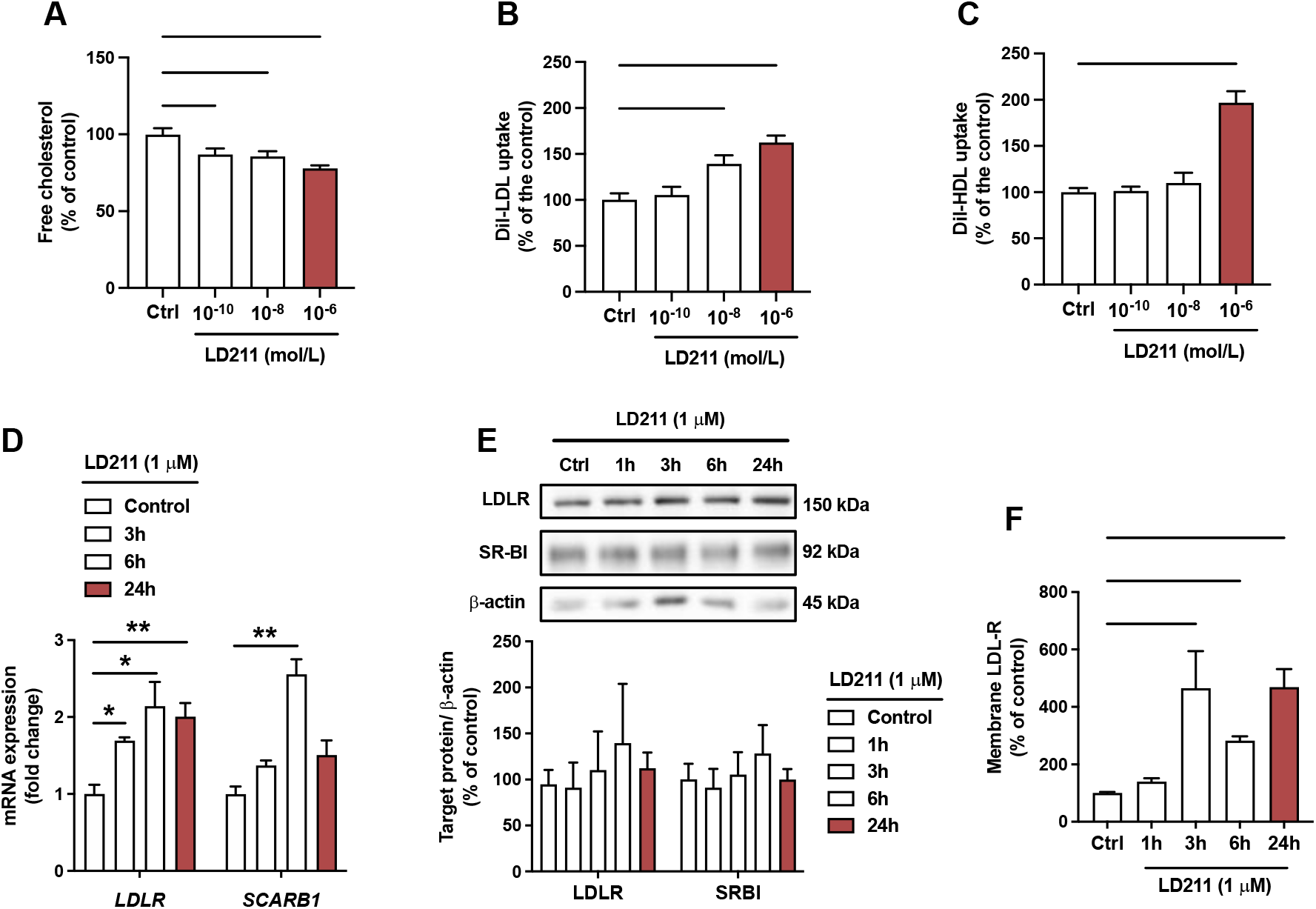
Selective activation of MC1-R mimics the actions of α-MSH in HepG2 cells. **(A)** Quantification of free cholesterol content using filipin staining in HepG2 cells treated with different concentrations of the selective MC1-R agonist LD211 (0.1 nM, 10 nM or 1 μM) for 24 hours. **(B, C)** Quantification of LDL and HDL uptake in HepG2 cells treated with different concentrations (0.1 nM, 10 nM or 1 μM) of LD211 for 24 hours. **(D)** qPCR analysis of *LDLR* and *SCARB1* expression in HepG2 cells treated with 1 μM LD211 for 3, 6 or 24 hours. **(E)** Representative Western blots and quantification of LDL-R and SR-BI proteins levels in HepG2 cells treated with 1μM LD211 for 1, 3, 6 or 24 hours. (**F**) Quantification of cell surface LDLR by flow cytometry in HepG2 cells treated with 1μM LD211 for 24 hours. Values are mean ± SEM, \**P<0*.*05*, \*\**P<0*.*01*, \*\*\**P<0*.*001* and \*\*\*\**P<0*.*0001* for the indicated comparisons by one-way ANOVA and Dunnet *post hoc* tests.

### MC1-R activation engages multiple signaling mechanisms to regulate cholesterol metabolism in HepG2 cells

We next aimed to investigate intracellular signaling cascades that might be activated in response to MC1-R stimulation. Since most MC-Rs are known to be coupled to Gs proteins, we first measured intracellular cAMP levels in α-MSH-treated HepG2 cells. However, α-MSH did not either increase or decrease cAMP levels, while the adenylyl cyclase activator forskolin, as a positive control, induced a robust increase in cAMP level (**Figure 7A**). We also screened other potential signaling pathways of MC1-R and found that mitogen-activated protein kinases, ERK (extracellular-signal-regulated kinase) and JNK (c-Jun N-terminal kinase), are affected by α-MSH treatment. The highest concentration of α-MSH (1 μM) reduced ERK phosphorylation (p-ERK1/2) at 5- and 15-min time points (**Supplementary Figure 6A**). Profiling of the concentration-response however revealed that the phosphorylation levels of ERK and JNK are most significantly reduced at the lowest tested concentration (0.1 nM) of α-MSH and the effects tend to fade away towards higher concentrations (**Figure 7B, C**). In addition, α-MSH induced a rapid phosphorylation (at 5 min) of AMP-activated protein kinase (AMPK) (**Figure 7D**) and the maximal effect was observed with 1 μM concentration (**Supplementary Figure 6C**), indicating a more conventional concentration-response. Treatment of HepG2 cells with α-MSH also progressively reduced the phosphorylation of protein kinase B (Akt) (**Supplementary Figure 6D**). The reduction in p-Akt level appeared to be driven by AMPK phosphorylation, since the AMPK inhibitor dorsomorphin increased Akt phosphorylation and abolished the effect of α-MSH on p-Akt level (**Supplementary Figure 6E**). Finally, we tested whether the α-MSH-induced reduction in cellular cholesterol content is mediated by AMPK phosphorylation. Interestingly, the AMPK inhibitor dorsomorphin increased cholesterol content and reversed the concentration-response to α-MSH (**Figure 7E**). In the presence of dorsomorphin, the lowest concentration of α-MSH showed the strongest response, while the effect of 1 μM α-MSH was completely blocked (**Figure 7E**).

**Figure 7.**
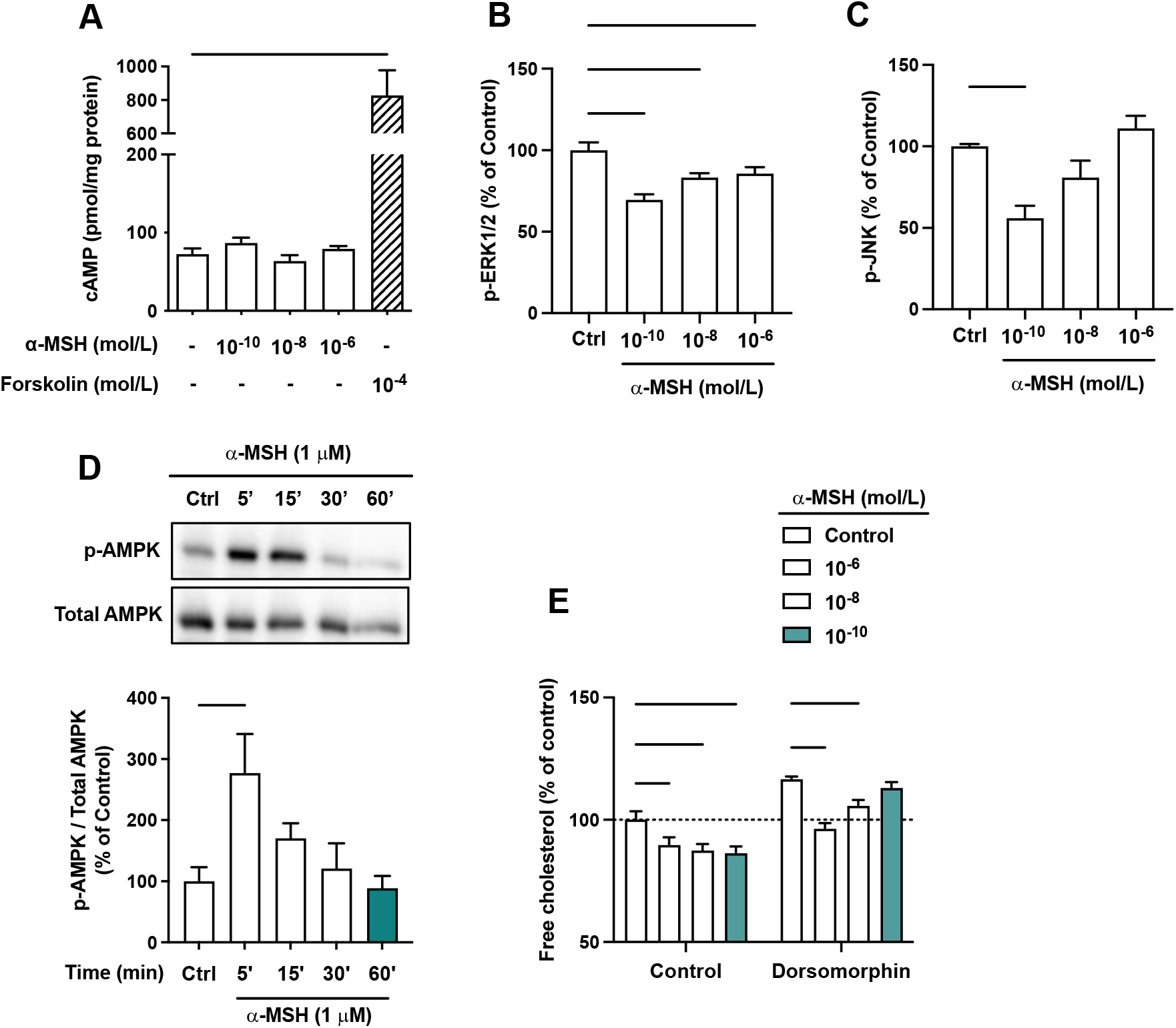
The effects of α-MSH on intracellular signaling pathways in HepG2 cells. **(A)** Quantification of intracellular cAMP level in HepG2 cells treated with different concentrations of α-MSH (0.1 nM, 10 nM or 1 μM) for 30 min. The adenylyl cyclase activator forskolin (10 μM) was used as a positive control. **(B, C)** Quantification of phosphorylated ERK1/2 and JNK by ELISA assays in HepG2 cells treated with different concentrations of α-MSH (0.1 nM, 10 nM or 1 μM) for 10 min. **(D)** Representative Western blots and quantification of phosphorylated AMPK level (p-AMPK normalized against total AMPK) in HepG2 cells treated with 1 μM α-MSH for 5, 15, 30 or 60 minutes. (**E**) Quantification of free cholesterol content using filipin staining in HepG2 cells treated with different concentrations of α-MSH (0.1 nM, 10 nM or 1 μM) for 24 hours in the presence or absence of the AMPK inhibitor dorsomorphin (1 μM). Values are mean ± SEM, \**P<0*.*05*, \*\**P<0*.*01* and \*\*\*\**P<0*.*0001* for the indicated comparisons by one-way ANOVA and Dunnet *post hoc* tests.

To verify the dependence of the observed effects on MC1-R activation, we repeated the signaling experiments using LD211 as a selective MC1-R agonist. Corroborating the findings from α-MSH-treated cells, LD211 had no effect on cAMP level but reduced the phosphorylation level of ERK1/2 and JNK most notably at the lowest tested concentration (**Supplementary Figure 7A-C)**. LD211 also induced phosphorylation of AMPK and reduced the level of p-Akt (**Supplementary Figure 7D, E)**, thus closely matching the phenotype of α-MSH-treated cells. Collectively, the results demonstrate that α-MSH evokes multiple signaling pathways and that the effects of α-MSH on cholesterol metabolism are not reliant on one single pathway.

## Discussion

In the present study, we investigated the role of MC1-R in hepatocytes and its possible involvement in cholesterol and bile acid metabolism. Firstly, extending previous observations on the hepatic expression of *Mc1r* mRNA, we show that MC1-R protein is widely present in the mouse liver and downregulated in response to feeding a cholesterol-rich diet. Secondly, hepatocyte-specific MC1-R deficiency rendered mice susceptible for enhanced accumulation of cholesterol and triglycerides in the plasma and liver. Loss of MC1-R signaling in hepatocytes disturbed also bile acid metabolism. Thirdly, *in vitro* experiments using HepG2 cells revealed that triggering MC1-R signaling either with the endogenous ligand α-MSH or the selective MC1-R agonist LD211 reduced cellular cholesterol content and enhanced the uptake of HDL and LDL cholesterol, which are preventive mechanisms against hypercholesteremia and associated cardiovascular complications.

Previous findings of *MC1R* mRNA expression in the rat and human liver (Gatti et al., 2006; López et al., 2007; Malik et al., 2012) and that global MC1-R deficiency aggravated hypercholesterolemia in Apoe^-/-^ mice (Rinne et al., 2018) led us to hypothesize that MC1-R regulates cholesterol metabolism in hepatocytes. *Mc1r* mRNA expression in the rat liver was previously found to increase after turpentine oil-induced acute phase response (Malik et al., 2012), while *MC1R* was downregulated in liver biopsies from brain-dead organ donors (Gatti et al., 2006), suggesting that inflammatory processes might modulate hepatic MC1-R expression. Here we show that the MC1-R is present also in the mouse liver that its mRNA and protein levels are reduced after feeding mice cholesterol-rich Western diet. We also analyzed MC1-R expression in HepG2 cells to gain further insight into the regulation of MC1-R expression in hepatocytes. Intriguingly, acute changes in cellular cholesterol load, as produced by treatment with atorvastatin or LDL-cholesterol, did not affect MC1-R protein level, while treatment with palmitic acid evoked an immediate reduction in MC1-R expression. These findings suggest that cholesterol *per se* does not regulate MC1-R expression but it might be rather modulated by inflammatory processes that are triggered by lipid overload.

To investigate the regulatory role of hepatic MC1-R, we generated *L-Mc1r*^*-/-*^ mice and found that hepatocyte-specific MC1-R deficiency induced hypercholesteremia and higher liver weight, which was accompanied by increased total cholesterol and triglyceride levels in the liver. *L-Mc1r*^*-/-*^ mice closely phenocopied the features of Apoe^-/-^ mice with global deficiency of MC1-R (Rinne et al., 2018), which displayed enhanced hypercholesteremia and cholesterol accumulation in the liver. Thus, the present findings suggest that the disturbed cholesterol metabolism previously observed in global MC1-R deficient mice was attributable to loss of MC1-R signaling in hepatocytes. The precise mechanism leading to enhanced hepatic lipid accumulation and hypercholesterolemia in *L-Mc1r*^*-/-*^ mice is not yet clear. Gene expression analysis in the liver showed upregulation of *Srebp2* and its target genes *Hmgcr, Dhcr7, Ldlr* and *Pcsk9*. Under normal physiological conditions, SREBP-1c preferentially activates the genes of fatty acid and triglyceride biosynthesis pathway, while SREBP-2 is considered as the master regulator of cholesterol metabolism (Hua et al., 1993; Yokoyama et al., 1993). Accordingly, the finding of increased *Srebp2* expression and the lack of effect on *Sreb1c* expression in *L-Mc1r*^*-/-*^ mice implies that hepatic MC1-R signaling predominantly regulates cholesterol rather than fatty acid metabolism. Considering that both HMGCR and DHCR7 are crucially involved in the cholesterol biosynthetic pathway (Prabhu et al., 2014; Shi et al., 2022), the upregulation of *Hmgcr* and *Dhcr7* is likely to enhance cholesterol synthesis and could thus explain in part the propensity for cholesterol accumulation in the liver of *L-Mc1r*^*-/-*^ mice.

In the presence of excess cellular cholesterol, transcriptional induction and posttranslational activation of SREBP-2 should be attenuated, which in turn downregulates *Hmgcr* and *Dhcr7* and reduces cholesterol synthesis as a counterregulatory mechanism. Therefore, given the increase in hepatic cholesterol content, it was unexpected that *Srebp2* expression was upregulated in the liver of *L-Mc1r*^*-/-*^ mice. These data could indicate that hepatic MC1-R signaling directly controls the transcription of *Srebp2* or its activation through proteolytic processing and translocation to the nucleus, which turns on gene expression for cholesterol synthesis. Alternatively, loss of MC1-R signaling might disturb cholesterol delivery to the endoplasmic reticulum (ER), which is sensed as cholesterol depletion in the cells and activates proteolytic release of SREBP2 from the ER and subsequent translocation to the nucleus. However, further research is warranted to dissect the mechanism by which hepatic MC-1R signaling regulates *Srebp2* expression and the downstream gene cluster for cholesterol biosynthesis.

A central feature of *L-Mc1r*^*-/-*^ mouse model was liver steatosis that occurred on a normal chow diet and without any signs of increased susceptibility for obesity. Although the precise mechanisms leading to NAFLD remain unclear, human and animal studies have shown that under normal caloric intake, the development of NAFLD is strongly associated with disturbance in liver cholesterol metabolism (Kainuma et al., 2006; Min et al., 2012; Simonen et al., 2011). Hence, the increased levels of plasma and liver cholesterol in *L-Mc1r*^*-/-*^ mice may have secondarily led to accumulation of triglycerides. Simple steatosis can progress to nonalcoholic steatohepatitis (NASH) that is characterized by progressive inflammation, oxidative stress and fibrosis. *L-Mc1r*^*-/-*^ mice showed enhanced liver fibrosis without any signs of elevated inflammation. Persistent lipid overload and consequent lipotoxicity could be the cause of fibrosis in *L-Mc1r*^*-/-*^ mice. However, MC1-R signaling has been shown to mediate anti-fibrotic effects and protect against skin fibrosis and systemic sclerosis (Böhm & Stegemann, 2014; Kondo et al., 2022), which opens the possibility that hepatocyte-specific MC1-R deficiency directly induces fibrogenesis.

*L-Mc1r*^*-/-*^ mice also displayed a unique bile acid profile with reduced secondary bile acid levels, particularly in the plasma and feces. Furthermore, the ratio of CA to CDCA was reduced in the plasma of *L-Mc1r*^*-/-*^ mice. This phenotype recapitulates some of the key features observed in mice with global deficiency of MC1-R (Rinne et al., 2018). Specifically, the relative amount of CA and the ratio of CA to CDCA were reduced in both mouse models. In the quest for possible explanation for this phenotype, we found that CYP8B1 was upregulated both at the mRNA and protein level in the liver of *L-Mc1r*^*-/-*^ mice. This contradicts the finding of reduced CA:CDCA ratio, which is predominantly determined by the enzymatic activity of CYP8B1 that is required for CA synthesis (Chiang, 2004). Therefore, it is plausible that BA synthesis *via* the classical pathway is disturbed in *L-Mc1r*^*-/-*^ mice due to dysfunctional CYP8B1, which leads to a compensatory enhancement of BA synthesis *via* the alternative pathway. This could explain the upregulation of StAR and CYP27A1, which operate in the mitochondria to feed the alternative BA pathway. Under physiological conditions, the majority of BAs are produced by the classical pathway, while in liver diseases such as NAFLD, the alternative BA pathway may become more dominant when compensating disturbances in the classical BA synthesis pathway (Chiang, 2004; Crosignani et al., 2007; Lake et al., 2013). Patients with NASH and NASH-driven hepatocellular carcinoma have also been reported to have increased hepatic StAR expression (F. Caballero et al., 2009; Conde de la Rosa et al., 2021).

In terms of BA transport, the upregulation of NTCP and downregulation of MRP4 in *L-Mc1r*^*-/-*^ mice might also indicate compensatory changes to disrupted BA synthesis. Increased NTCP is likely to enhance BA uptake from the portal circulation and thus enterohepatic circulation of BAs, while reduced MRP4 expression prevents excessive spillover of BAs into systemic circulation. These changes synergistically help to maintain the liver BA pool in the presence of disturbed BA synthesis. Reduced MRP4 expression in *L-Mc1r*^*-/-*^ mice also provides mechanistic explanation for the finding of reduced BA levels in the plasma. It remains however to be determined whether the dysregulated BA metabolism contributes to the hypercholesteremia and increased hepatic lipid accumulation in *L-Mc1r*^*-/-*^ mice. For instance, CYP7A1 deficiency in both humans and mice causes hypercholesterolemic phenotype with increased hepatic cholesterol content (Erickson et al., 2003; Pullinger et al., 2002). Because there is reciprocal interaction between fatty liver disease and dysregulated BA metabolism, it is difficult to determine whether MC1-R deficiency *per se* disturbs BA metabolism or whether enhanced lipid accumulation in *L-Mc1r*^*-/-*^ mice causes a defect in BA metabolism. However, *in vitro* experiments with HepG2 cells support the notion that hepatic MC1-R signaling might directly affect BA metabolism, since treatment with the endogenous MC1-R agonist α-MSH modestly increased CYP8B1 expression, CA synthesis and CA:CDCA ratio.

In good agreement with our findings in *L-Mc1r*^*-/-*^ mice, *in vitro* experiments using HepG2 cells demonstrated that triggering MC1-R signaling with the endogenous agonist α-MSH or the synthetic agonist LD211 induced a reverse phenotype, namely reduction in cellular cholesterol content. MC1-R activation was also associated with enhanced LDL and HDL uptake in HepG2 cells. Inhibition of cholesterol synthesis (*e*.*g*. by statins) is known to similarly reduce cellular cholesterol level in hepatocytes, which, in turn, upregulates LDL-R and reduces plasma total and LDL cholesterol. In the case of MC1-R activation, the increase in LDL-R expression and LDL uptake might be partly independent of the effect on cellular cholesterol content, since the upregulation of LDL-R occurred rapidly after α-MSH treatment (at 1-hour time point) and before any noticeable change in cellular cholesterol level. Therefore, the data suggest that the upregulation of LDL-R as well as SR-BI is attributable mainly to direct transcriptional induction by MC1-R signaling. Supporting the therapeutic relevance of this finding, we had previously observed that chronic treatment of atherosclerotic mice with a selective MC-R agonist increased LDL-R expression in the liver and reduced plasma total cholesterol concentration (Rinne et al., 2017). The underlying mechanism remained obscure in that study, but the present findings suggest that the upregulation of LDL-R expression was a consequence of MC1-R activation in hepatocytes. In the current study, we also found that MC1-R activation reduced cellular cholesterol content in HepG2 cells but we were unable to pinpoint the exact molecular-level mechanism for this effect. Since we did not observe any change in the expression of the major cholesterol biosynthetic enzymes (HMGCR or DHCR7), we speculate that MC1-R signaling might inhibit other enzymes in the cholesterol biosynthesis pathway or modulate the phosphorylation state of HMGCR that determines its catalytic activity (Burg & Espenshade, 2011). Alternatively, or in addition to other mechanisms, enhanced cholesterol turnover into BAs, as evidenced by increased CA production and CYP8B1 expression in α-MSH-treated HepG2 cells, might reduce cellular cholesterol content. In any case, these data uncover a functional role for MC1-R signaling in hepatic cholesterol metabolism that might be of therapeutic relevance in the management of hypercholesterolemia.

In terms of intracellular signaling, MC1-R activation in HepG2 cells evoked phosphorylation of AMPK and inhibition of ERK1/2 and JNK without any effect on cAMP levels. Melanocortin receptors are classically coupled to Gs protein and cAMP-dependent signaling (Rodrigues et al., 2015), but the present findings demonstrate that the MC1-R-mediated effects on cholesterol metabolism occur in a cAMP-independent manner. Mechanistic experiments further revealed that the reduction of cellular cholesterol level by α-MSH was partially reversed after AMPK inhibition with dorsomorphine. However, in the presence of dorsomorphine, low concentrations of α-MSH were still effective in reducing cellular cholesterol content and the concentration-response closely mirrored the profiles observed in p-JNK and p-ERK1/2 levels in α-MSH-treated HepG2 cells (*i*.*e*. U-shaped concentration-response). These results suggest that the α-MSH-mediated reduction in cellular cholesterol content relies on multiple pathways involving AMPK and MAPK signaling pathways. Early *in vitro* studies have established that AMPK activation reduces cholesterol synthesis by inducing an inactivating phosphorylation of HMGCR (Clarke & Hardie, 1990; Steinberg & Kemp, 2009), while inhibition of AMPK increases cholesterol synthesis and cholesterol accumulation in the liver (Loh et al., 2018). The link between MAPK signaling and cholesterol metabolism has not been widely studied but *in vitro* experiments using HepG2 cells have demonstrated that ERK1/2 activation phosphorylates SREBP2 (Kotzka et al., 2004), which, in turn, is likely to induce e.g. *HMGCR* transcription. The role of hepatic JNK signaling is even less clear in this regard, but some evidence demonstrates that it contributes to diet-induced obesity and hepatic steatosis as well as to cholesterol and BA metabolism (Manieri et al., 2020; Manieri & Sabio, 2015; Vernia et al., 2014). Against this background, it is plausible that both AMPK phosphorylation and inhibition of ERK1/2 and JNK signaling are involved in mediating the effects of α-MSH on cholesterol metabolism in hepatocytes. However, further experiments are warranted to dissect the exact signaling mechanism(s) of MC1-R activation that regulate cholesterol metabolism, *e*.*g*., to determine which G-protein subtype is involved in this regulation.

In conclusion, our study uncovers a novel role for MC1-R signaling in hepatic cholesterol and BA metabolism. Hepatocyte-specific MC1-R deficiency increased plasma cholesterol and TG concentration, disturbed BA metabolism and led to signs of hepatic steatosis and fibrosis. Conversely, MC1-R activation in hepatocytes reduced cellular cholesterol content and increased LDL and HDL uptake, which are preventive mechanisms against hypercholesterolemia and the progression of NAFLD.

## Materials and methods

### Mice

All experiments were performed on adult (3-6 months) female mice. Mice were housed in groups of littermates on a 12 h light/dark cycle. The numbers of mice studied in each experiment are given in the figure legends. Mice were maintained on a regular chow diet (# 2916C, Teklad Global diet, Envigo) for the entire experimental duration unless otherwise stated. In each experiment, mice were euthanized *via* CO_2_ asphyxiation, blood was withdrawn, and whole liver was excised and weighed. The experiments were approved by the local ethics committee (Animal Experiment Board in Finland, License Numbers: ESAVI/6280/04.10.07.2016 and ESAVI/1260/2020) and conducted in accordance with the institutional and national guidelines for the care and use of laboratory animals.

Eight-week-old C57BI/6J mice (Janvier Labs, France) were fed a regular chow diet or Western-type diet (RD Western Diet, D12079B, Research Diets Inc, NJ, USA) for 12 weeks and used for the quantification of MC1-R mRNA and protein levels in the liver. In addition, hepatocyte specific MC1-R knock-out mice (*L-Mc1r*^*-/-*^) were generated by breeding mice homozygous for a floxed Mc1r allele (*Mc1r*^*fl/fl*^ mice, the Jackson Laboratory, strain #029239) (Takeo et al., 2016) with transgenic Alb-Cre^+/-^ mice (B6N.Cg-Speer6-ps1Tg(Alb-cre)21Mgn/J, the Jackson Laboratory, strain #018961) (Postic et al., 1999). Age-matched Mc1r^fl/fl^ (Alb-Cre^-/-^) and Alb-Cre^+/-^ Mc1r^wt/fl^ (*L-Mc1r*^*+/-*^) mice were used as controls. Mice were weighed once a week during a monitoring period of 8 weeks (from 8 to 16 weeks of age). Body composition was determined at the start and end of the 8-week monitoring period by quantitative nuclear magnetic resonance (NMR) scanning (EchoMRI-700, Echo Medical Systems, Houston, TX, USA). At the end of the experiment, genomic DNA samples from the liver were genotyped for the recombined allele using the following primers: ACC ACT GCG TGC TAT CCT G (*Mc1r* 5’forward), ACC CCT TCC CTT GAG GAG T (*Mc1r* 5’ reverse) and GAA CTC TGA GGT CAC TAT TTT CTG GAG A (*Mc1r* 3’ reverse).

### Cell culture

The HepG2 cell line was purchased from ATCC (American Type Culture Collection, Rockville, MD, USA; HB-8065) and maintained in DMEM (Dulbecco’s modified Eagle’s medium; Sigma-Aldrich) supplemented with 10 % (v/v) heat-inactivated fetal bovine serum (FBS; Gibco), 100 U/ml penicillin (Gibco), 100 μg/ml streptomycin (Gibco) at 37°C in a humid atmosphere with 5 % CO_2_. To study the regulation of MC1-R expression, HepG2 cells were serum-deprived (0.5 % FBS) for 16 h and thereafter treated with 200 μg/ml LDL (CliniSciences), 10 μM atorvastatin (Sigma-Aldrich) or 500 μM palmitic acid (Sigma-Aldrich) for 1, 3, 6 or 24 h. To study the effects of melanocortin system activation, cells were seeded on 12- or 24-well plates and treated with the non-selective MC-R agonist α-MSH (abcam, # ab120189) or the selective MC1-R agonist LD211 as indicated in the figure legends (Doedens et al., 2010) (compound 28 in the original publication).

### Histology and immunohistochemistry

A transverse piece of the left lobe was fixed in 10% formalin overnight followed by embedding in paraffin. Four μm-thick serial sections were stained with hematoxylin and eosin (H&E), Picrosirius Red (abcam, # ab150681) or used for MC1-R immunohistochemistry as previously described (Rinne et al., 2015, 2017). Briefly, sections were incubated in 10 mM sodium citrate buffer (pH 6) for 20 min in a pressure cooker for antigen retrieval. Thereafter, sections were quenched in 1 % H_2_O_2_ for 10 min and blocked in 5 % normal horse serum containing 1 % BSA. After blocking, sections were incubated overnight with a primary antibody against MC1-R (Alomone Labs, Jerusalem, Israel, # AMR-025) followed by biotinylated horseradish peroxidase-conjugated secondary antibody incubation and detection with diaminobenzidine (ABC kit, Vector Labs, Burlingame, USA). For isotype control, a consecutive heart section was treated similarly except that the primary MC1-R antibody was replaced by purified normal rabbit IgG (Novus Biologicals, Littleton, CO, USA, # NB810-56910). Sections were counterstained with hematoxylin (CarlRoth), cover-slipped and then scanned with Pannoramic 250 or Pannoramic Midi digital slide scanner (3DHISTECH Kft, Budapest, Hungary). To visualize hepatic lipid content, a transverse piece of the left liver lobe was embedded in O.C.T. compound (Tissue-Tek), cryosectioned and stained with Oil Red O.

### RNA isolation, cDNA synthesis and quantitative RT-PCR

HepG2 cell samples were collected into QIAzol Lysis Reagent and total RNA was extracted using Direct-zol RNA Miniprep (Zymo Research, CA, USA). Liver samples were first homogenized in QIAzol Lysis Reagent (Qiagen) using the Qiagen TissueLyser LT Bead Mill (QIAGEN, Venlo, Netherlands). Total RNA from each sample was extracted and reverse-transcribed to cDNA with PrimeScript RT reagent kit (Takara Clontech) according to the manufacturer’s instructions. The RNA quality and concentration was evaluated by Nanodrop. Quantitative real-time polymerase chain reaction (RT-PCR) was performed using SYBR Green protocols (Kapa Biosystems, MA, USA) on a real-time PCR detection system (Applied Biosystems 7300 Real-Time PCR system). Each sample was run in duplicate. Target gene expression was normalized to the geometric mean of two housekeeping genes (β-actin and ribosomal protein S29 or GAPDH) using the delta-Ct method and results are presented as relative transcript levels (2^-ΔΔCt^). Primer sequences are presented in **Supplementary Tables I** and **II**.

### Immunoblotting

Liver and HepG2 samples were lysed in RIPA buffer (50 mM NaCl, 1 % Triton X-100, 0.5 % Sodium deoxycholate, 0.1 % SDS, pH 8.0) supplemented with protease and phosphatase inhibitor cocktail (ThermoFisher, #A32961). Liver samples were additionally homogenized using the Qiagen TissueLyser LT Bead. Equal amounts (30 μg) of total protein were separated by 10 % SDS-polyacrylamide gel electrophoresis (SDS-PAGE) and transferred onto nitrocellulose membranes (GE Healthcare). After blocking with Tris-Buffered Saline (Sigma-Aldrich) containing 0.1 % Tween^®^ 20 detergent (Sigma-Aldrich) and 5 % skimmed milk (Carl Roth) for 1 h at room temperature (RT), membranes were incubated with specific primary antibodies for MC1-R (Alomone Labs, #AMR-025), LDLR (NovusBio, Littleton, CO, USA, #NBP1-06709), SR-BI (NovusBio, #NB400-104), HMGCR (NovusBio, #NBP2-66888), DHCR7 (abcam, #ab103296), MRP4 (Cell Signaling Tech, Frankfurt, DE, #12857), StAR (Cell Signaling Tech, #8449), CYP8B1 (St John’s Laboratory Ltd, #STJ92607), phospho-AMPKα (Cell Signaling Tech, #2535), AMPKα (Cell Signaling Tech, #2532), phospho-Akt (R&D Systems, #AF887) and Akt (R&D Systems, #MAB2055) over-night at + 4°C. Next day, membranes were washed and incubated with Horseradish peroxidase (HRP)-conjugated secondary antibodies (Cell Signaling Tech) for 1 h at RT. Proteins were visualized using a chemiluminescence (ECL) kit (Millipore, MA, USA). Target protein expression was normalized to β-actin (Sigma-Aldrich, #2066) or vinculin (Bio-Rad, #MCA465GA) to correct for loading and band densities were analyzed using ImageJ software (NIH, Bethesda, MD, USA).

### Plasma and liver extract analyses

Plasma samples were obtained from EDTA-anticoagulated whole blood after centrifugation. Plasma total cholesterol and triglycerides concentrations were determined using enzymatic colorimetric assays (CHOD-PAP and GPO-PAP, mtiDiagnostics, Idstein, Germany) according to the manufacturer’s protocols. For the determination of hepatic lipid content, liver samples (∼100 mg) were homogenized in 500 μl of PBS with 0.1% NP-40 (Abcam) using TissueLyser and then centrifuged to remove insoluble components (Nuutinen et al., 2018; Rinne et al., 2018). Cholesterol and triglycerides concentrations were quantified in the liver homogenates using CHOD-PAP and GPO-PAP reagents.

### Bile acid measurements

Bile acids were measured in plasma, liver and fecal samples. Feces were collected over 48 h from individually housed mice. Total bile acids were extracted and analyzed by an ultra high-performance liquid chromatography tandem mass spectrometry method (UHPLC-MS/MS) as previously described (Jäntti et al., 2014).

### Filipin staining of cellular free cholesterol

HepG2 cells were seeded (40 000 cells/well) on 96-well plates (PhenoPlate™, PerkinElmer) and grown until the cells reached 70 % confluency. Thereafter, cells were first serum-deprived (0.5 % FBS) for 16 h and then treated with either α-MSH or LD211 for 24 h. After the treatment, cells were washed with PBS and fixed with 4 % paraformaldehyde (Sigma-Aldrich) for 15 min at room temperature. Cells were subsequently washed with PBS and incubated with 1.5 mg/ml glycine (Sigma-Aldrich, 10 min at RT) to quench unreacted paraformaldehyde followed by washing with PBS. Cells were stained with 50 ug/ml of Filipin (Sigma-Aldrich, #F9765) for 1 h at 37°C and washed with PBS. Fluorescence signal was measured with EnSight Multimode Plate Reader (PerkinElmer) with the 360 nm excitation and 480 emission wavelengths.

### LDL and HDL uptake assay

HepG2 cells were seeded (40 000 cells/well) on 96-well plates (PhenoPlate™, PerkinElmer), serum-deprived (0.5 % FBS) for 16h and then treated with α-MSH or LD211 for 18 h at 37°C. After the treatment, cells were washed with PBS and incubated with fluorescently-labeled HDL (Dil-HDL, 20 μg/ml, CliniSciences) or LDL (Dil-LDL, 10 μg/ml, CliniSciences) for 4h at 37°C. After the incubation, cells were again washed with PBS and fluorescence signal was measured with EnSight Multimode Plate Reader (PerkinElmer) with the 549 nm excitation and 565 emission wavelengths and normalized against cell confluency.

### Flow cytometric analysis of cell surface LDLR expression

HepG2 cell were treated as indicated in the figure legends, washed with PBS and detached using EDTA. To quantify the expression of LDLR on the cell surface, HepG2 cells were stained with PE-conjugated anti-human LDLR antibody (clone C7, BD Biosciences) and then analyzed with LSR Fortessa (BD Biosciences) and FlowJo software (FlowJo, LLC, Ashland, USA).

### Cyclic AMP determination

To measure intracellular cAMP concentrations, HepG2 cells were pretreated with 3-isobutyl-1-methylxanthine (0.1 mM, IBMX, Sigma-Aldrich) for 30 min and then stimulated with α-MSH or the selective MC1-R agonist LD211 (0.1 nM, 10 nM or 1 μM) for 30 minutes. Cells were thereafter lysed with 0.1 M HCl and assayed for cAMP levels with a commercial kit (Cyclic AMP Select ELISA kit, Cayman Chemical, #501040) according to manufacturer’s instructions. Results were normalized against total protein concentrations (Pierce™ BCA Protein Assay Kit, ThermoFisher) and expressed as percentage of control samples that were left untreated.

### Enzyme-linked immunosorbent assays (ELISA) of phosphorylated ERK1/2 and JNK

HepG2 cells were stimulated with α-MSH or the selective MC1-R agonist LD211 as indicated in the figure legends. Cells were thereafter lysed with Lysis Buffer #6 (R&D Systems) and assayed for the expression levels of phospho-ERK1 (T202/Y204)/ERK2 (T185/Y187) and phospho-JNK (T183/Y185 for JNK1/2 and T221/Y223 for JNK3) with commercial kits (DuoSet IC ELISA, R&D Systems, # DYC1018B and # DYC1387B) according to manufacturer’s instructions. Results were normalized against total protein concentrations (Pierce™ BCA Protein Assay Kit, ThermoFisher).

### Statistics

Statistical analyses were performed with GraphPad Prism 9 software (La Jolla, CA, USA). Statistical significance between the experimental groups were determined by two-tailed, unpaired Student’s t test or one-way or two-way ANOVA followed by Dunnet *post hoc* tests. The D’Agostino and Pearson omnibus normality test method was utilized to check the normality of the data. Possible outliers in the data sets were identified using the regression and outlier removal (ROUT) method of Q-level of 1 %. Data are expressed as mean ± standard error of the mean (SEM). Results were considered significant for P<0.05.

## Author contributions

K.T., E.S., and P.R. conceptualized the study and designed the study methodology. K.T., J.J.K., K.S., I.P., and P.R. performed experiments. M.C. synthetized selective MC1-R agonist. K.T., E.S. and P.R. analyzed data. M.C., E.S. and P.R. acquired funding. K.T. and P.R. wrote the manuscript. All authors provided critical feedback and approved the final version of the manuscript.

## Acknowledgements

We thank Sanna Bastman, Salla Juhantalo and Johanna Jukkala for their excellent technical support. Erica Nyman and Marja-Riitta Kajaala (Histology core facility, University of Turku) are acknowledged for processing histology samples. BA measurements were performed at the Turku Metabolomics Centre (Turku Bioscience Centre, Turku, Finland) with the support of Biocenter Finland. Turku Center for Disease Modeling, TCDM, Turku, Finland, (www.tcdm.fi), a member of Biocenter Finland, is acknowledged for setting up the genotyping protocol for *L-Mc1r*^*-/-*^ mice.

This work was supported by the Academy of Finland (grant 315351 to P.R.); the Sigrid Jusélius Foundation (to P.R.); the Finnish Cultural Foundation (to P.R.), the Finnish Foundation for Cardiovascular Research (to E.S. and P.R.); and the National Institutes of Health (grant GM-104080 to M.C.).

