## Supplementary Material for "Melanocortin 1 receptor regulates cholesterol and bile acid metabolism in the liver"

**Supplementary Table I. Quantitative RT-PCR primers for mouse genes.**

| **Gene name**<br>Accession number | **5'-3' primer sequence** |
|---|---|
| ***Abca1***<br>NM_013454.3 | Forward: gcagatcaagcatcccaact<br>Reverse: ccagagaatgtttcattgtcca |
| ***Abcg1***<br>NM_009593.2 | Forward: gggtctgaactgccctacct<br>Reverse: tactcccctgatgccacttc |
| ***Abcg5***<br>NM_031884.2 | Forward: tggatccaacacctctatgctaaa<br>Reverse: ggcaggttttctcgatgaactg |
| ***Abcg8***<br>NM_026180.3 | Forward: tgcccaccttccacatgtc<br>Reverse: atgaagccggcagtaaggtaga |
| ***Actb***<br>NM_007393.5 | Forward: tccatcatgaagtgtgacgt<br>Reverse: gagcaatgatcttgatcttca |
| ***Akr1d1***<br>NM_145364.2 | Forward: gaaaagatagcagaagggaaggt<br>Reverse: gggacatgctctgtattccataa |
| ***Bsep***<br>NM_021022.3 | Forward: aagctacatctgccttagacacagaa<br>Reverse: caatacaggtccgaccctctct |
| ***Ccl2***<br>NM_011333.3 | Forward: aggtccctgtcatgcttctg<br>Reverse: aaggcatcacagtccgagtc |
| ***Col1a1***<br>NM_007742.4 | Forward: gctcctcttaggggccact<br>Reverse: ccacgtctcaccattgggg |
| ***Cyp7a1***<br>NM_007824.2 | Forward: gatcctctgggcatctcaag<br>Reverse: agaggctgctttcattgctt |
| ***Cyp7b1***<br>NM_007825.4 | Forward: gaaaactcttcaaaggcaacatgg<br>Reverse: actggaaagggttcagaacaaatg |
| ***Cyp8b1***<br>NM_010012.3 | Forward: gccttcaagtatgatcggttcct<br>Reverse: gatcttcttgcccgacttgtaga |
| ***Cyp27a1***<br>NM_024264.5 | Forward: gcctcacctatgggatcttca<br>Reverse: tcaaagcctgacgcagatg |
| ***Dhcr7***<br>NM_007856.2 | Forward: gaggcgtccaagaaggtg<br>Reverse: gcagcccattcacctcatac |
| ***Fxr***<br>NM_001163700.1 | Forward: tccggacattcaaccatcac<br>Reverse: tcactgcacatcccagatctc |
| ***Hmgcr***<br>NM_008255.2 | Forward: tgattggagttggcaccat<br>Reverse: tggccaacactgacatgc |
| ***Hnf4a***<br>NM_008261.3 | Forward: accaagaggtccatggtgttt<br>Reverse: gtgccgagggacgatgtag |
| ***Hsd3b7***<br>NM_133943.2 | Forward: gggagctgcgtgtctttga<br>Reverse: gtggatggtctttggactggc |

| **Gene name** Accession number | **5'-3' primer sequence** |
|---|---|
| ***Il1b*** NM_008361.4 | Forward: tgtaatgaaagacggcacacc<br>Reverse: tcttctttgggtattgcttgg |
| ***Il6*** NM_031168.2 | Forward: ggccttccctacttcacaag<br>Reverse: atttccacgatttcccagag |
| ***Ldlr*** NM_010700.3 | Forward: gcgtaaagaggaggacactgtt<br>Reverse: ccaatctgtccagtacatgaagc |
| ***Lrh1*** NM_030676.3 | Forward: tgggaaggaagggacaatctt<br>Reverse: cgagactcaggaggttgttgaa |
| ***Lxra*** NM_013839.4 | Forward: tgggatgtccacgagtgactgttt<br>Reverse: tcccttaatgctacggaaggctct |
| ***Mrp3 (Abcc3)*** NM_029600.4 | Forward: ctgggtcccctgcatctac<br>Reverse: gccgtcttgagcctggataac |
| ***Mrp4 (Abcc4)*** NM_001163676.1 | Forward: ggcactccggttaagtaactc<br>Reverse: tgtcacttggtcgaatttgttca |
| ***Ntcp*** NM_011387.2 | Forward: gaagtccaaaaggccacactatgt<br>Reverse: acagccacagagagggagaaag |
| ***Osta (Slc51a)*** NM_145932.3 | Forward: aggcaggactcatatcaaacttg<br>Reverse: tgagggctatgtccactggg |
| ***Pcsk9*** NM_153565.2 | Forward: ttgcagcagctgggaactt<br>Reverse: ccgactgtgatgacctctgga |
| ***S29*** NM_009093.2 | Forward: atgggtcaccagcagctcta<br>Reverse: agcctatgtccttcgcgtact |
| ***Scarb1*** NM_016741.2 | Forward: gcccatcatctgccaact<br>Reverse: tcctgggagcccctttttact |
| ***Shp (Nr0b2)*** NM_011850.3 | Forward: tgggtcccaaggagtatgc<br>Reverse: gctccaagacttcacacagtg |
| ***Srebp1c*** NM_011480.4 | Forward: gatgtgcgaactggacacag<br>Reverse: catagggggcgtcaaacag |
| ***Srebp2*** NM_033218.1 | Forward: ccaaagaaggagagaggcgg<br>Reverse: cgccagacttgtgcatcttg |
| ***Stard1*** NM_011485.5 | Forward: atgttcctcgctacgttcaag<br>Reverse: cccagtgctctccagttgag |
| ***Tgfb1*** NM_011577.2 | Forward: ccgcaacaacgccatctatg<br>Reverse: cccgaatgtctgacgtattgaag |
| ***Tnf*** NM_013693.3 | Forward: ctgaacttcggggtgatcgg<br>Reverse: ggcttgtcactcgaattttgaga |

**Supplementary Table II. Quantitative RT-PCR primers for human genes.**

| **Gene name**<br>Accession number | **5'-3' primer sequence** |
|---|---|
| ***ACTB***<br>NM_001101.5 | Forward: caccattggcaatgagcggttc<br>Reverse: aggtctttgcggatgtccacgt |
| ***GAPDH***<br>NM_002046.7 | Forward: tcaaggctgagaacgggaag<br>Reverse: cgccccacttgattttggag |
| ***LDLR***<br>NM_000527.5 | Forward: ccacggtggagatagtgaca<br>Reverse: ctcacgctactgggcttctt |
| ***SCARB1***<br>NM_005505.5 | Forward: ctggcagaagcggtgact<br>Reverse: cagagcagttcatggggatt |

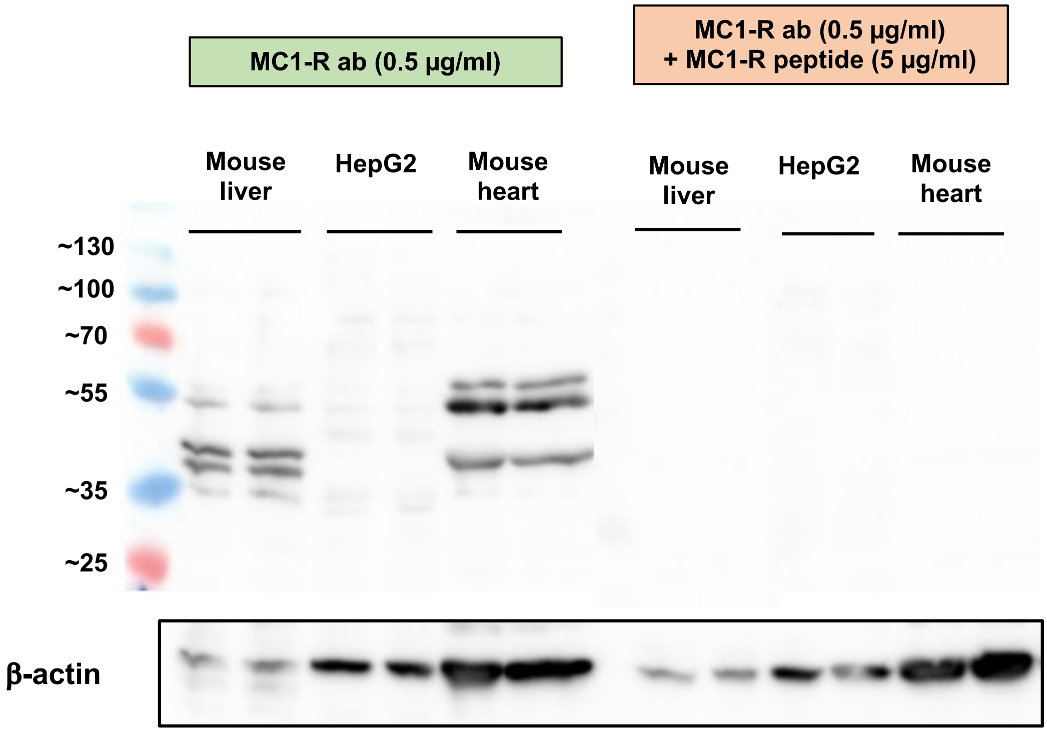


**Supplementary Figure 1.** Pre-adsorption control for MC1-R Western blotting. Western blot analysis of MC1-R protein expression in the mouse liver, HepG2 and mouse heart samples. The expression of β-actin is shown as loading control. Lanes on the right (same samples as on the left) were incubated in anti-MC1-R antibody solution that was premixed with a molar excess of a blocking MC1-R peptide.

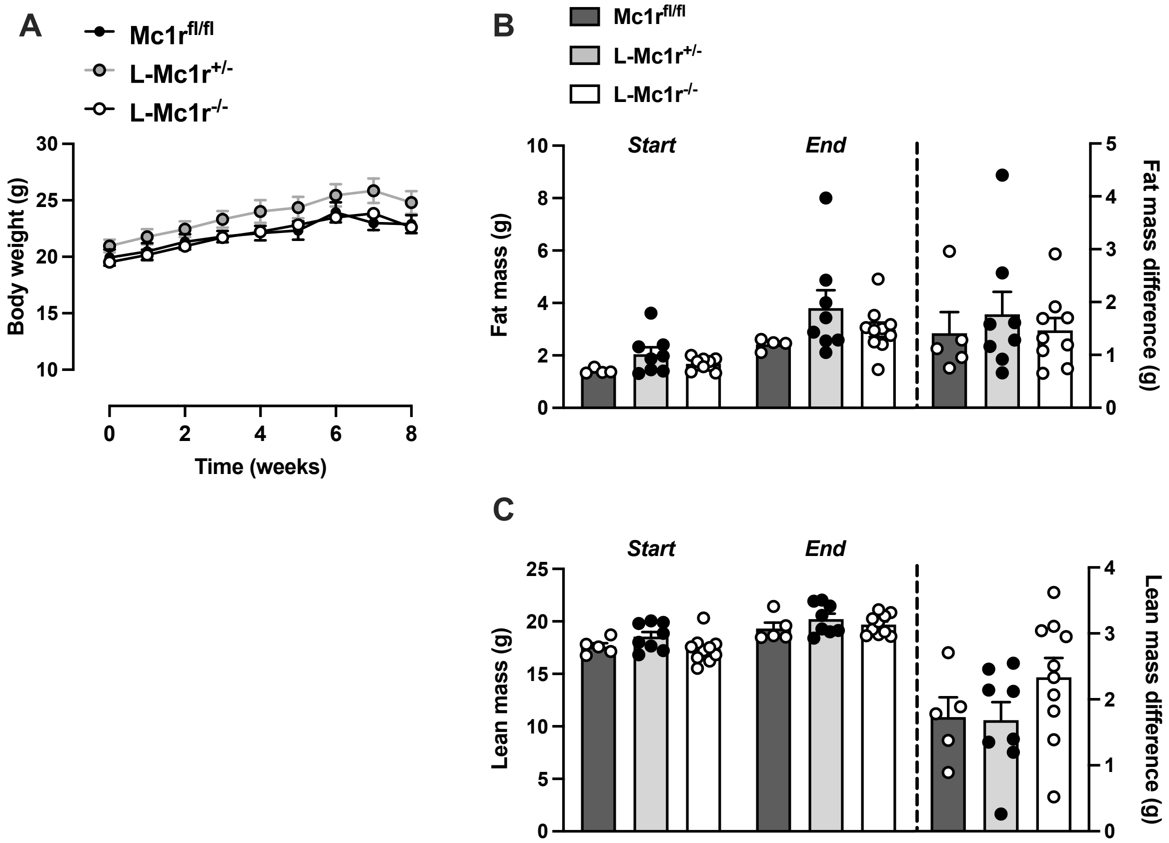


**Supplementary Figure 2.** Hepatocyte-specific MC1-R deficiency does not affect body weight or composition in chow-fed female mice. (**A**) Body weight curves of chow-fed *Mc1r*$^{fl/fl}$, *L-Mc1r*$^{+/-}$ and *L-Mc1r* mice. (**B** and **C**) Total fat and lean mass of chow-fed *Mc1r*$^{fl/fl}$, *L-Mc1r*$^{+/-}$ and *L-Mc1r* mice at the start and end of the body weight monitoring period. The change of fat and lean mass between the start and end of the experiment is also presented in the graphs. Values are mean ± SEM, mice n = 5-10 mice per group in each graph.

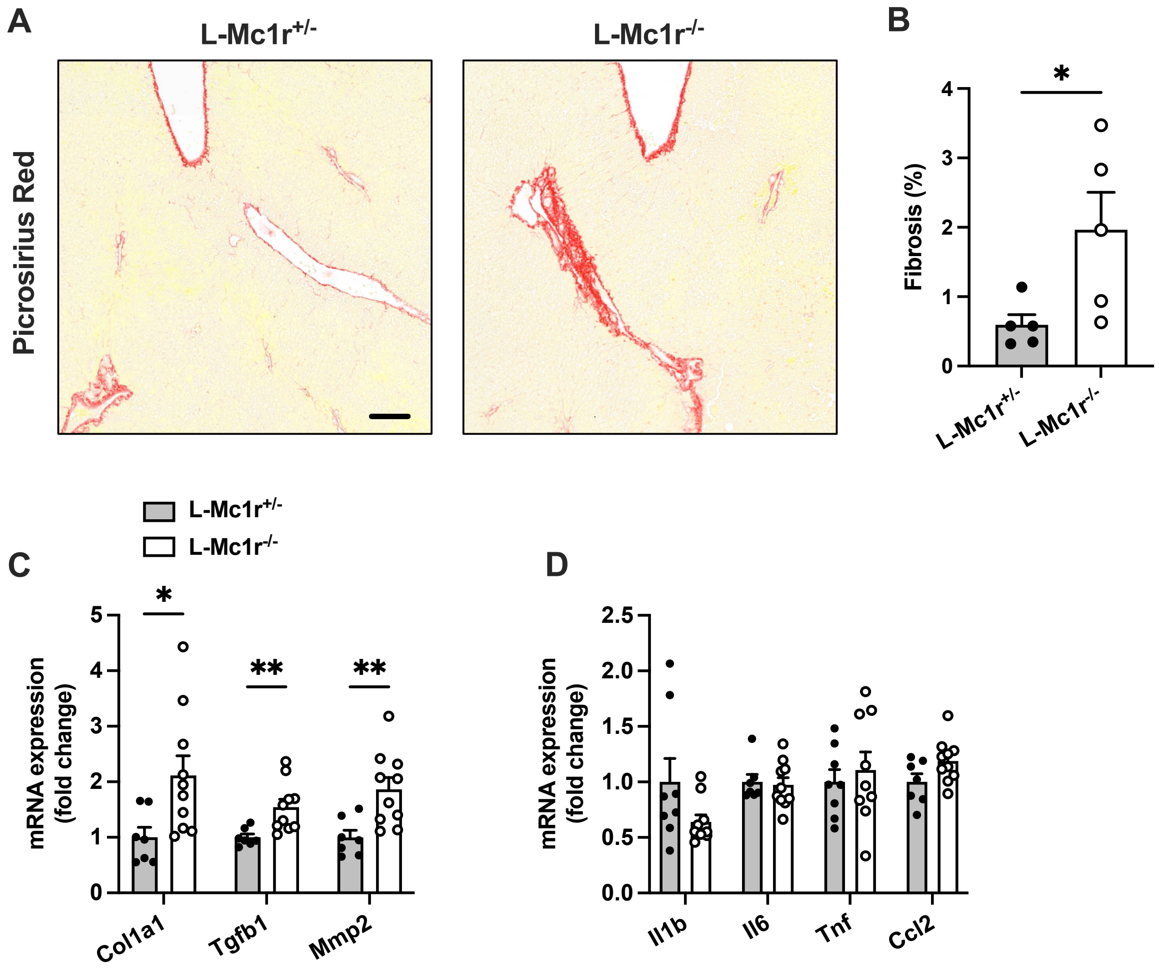


**Supplementary Figure 3.** Hepatocyte-specific MC1-R deficiency enhances liver fibrosis. (**A**) Representative Picrosirius Red-stained liver sections of chow-fed *L-Mc1r*$^{+/-}$ and *L-Mc1r* mice. Scale bar, 100 µm. (**B**) Quantification of fibrotic area (as percentage of total tissue area) in the liver of chow-fed *L-Mc1r*$^{+/-}$ and *L-Mc1r*$^{-/-}$ mice. n = 5 mice per group. qPCR analysis of fibrotic (**C**) and pro-inflammatory (**D**) genes in the liver of chow-fed *L-Mc1r*$^{+/-}$ and *L-Mc1r*$^{-/-}$ mice. Values are mean ± SEM, mice n = 8-10 mice per group in each graph. **P<0.05* and ***P<0.01* for the indicated comparisons by Student's t test. *Col1a1*, collagen, type I, alpha 1; *Tgfb1*, transforming growth factor beta 1; *Mmp2*, matrix metalloproteinase-2; *Il1b*, interleukin 1 beta; *Il6*, interleukin 6; *Tnf*, tumor necrosis factor; *Ccl2*, chemokine (C-C motif) ligand 2.

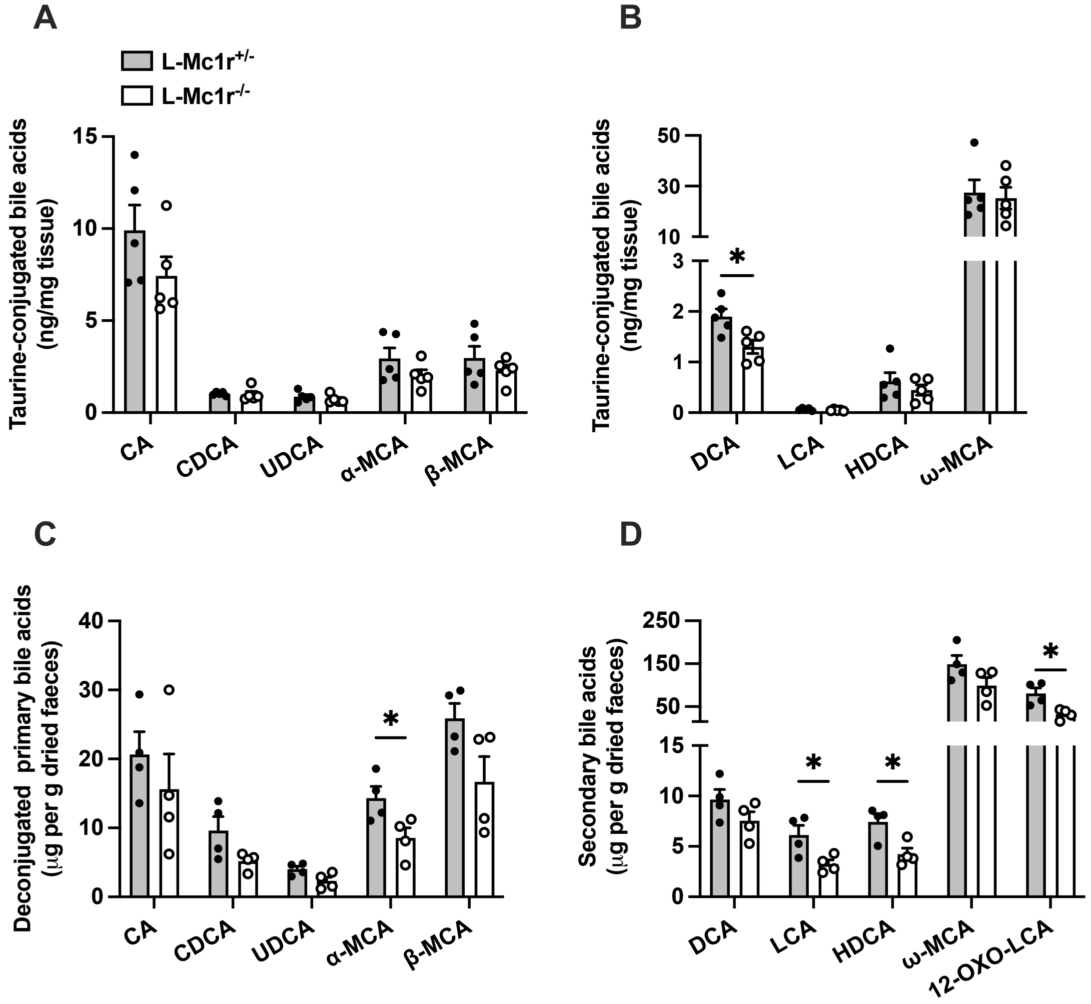


**Supplementary Figure 4.** Bile acid profiles in the liver and feces of *L-Mc1r*$^{-/-}$ mice. (**A**) Quantification of individual primary bile acids in the liver of chow-fed *L-Mc1r*$^{+/-}$ and *L-Mc1r*$^{-/-}$ mice. (**B**) Quantification of individual secondary bile acids in the liver of chow-fed *L-Mc1r*$^{+/-}$ and *L-Mc1r*$^{-/-}$ mice. (**C**) Quantification of individual primary bile acids in the feces of chow-fed *L-Mc1r*$^{+/-}$ and *L-Mc1r*$^{-/-}$ mice. (**D**) Quantification of individual secondary bile acids in the feces of chow-fed *L-Mc1r*$^{+/-}$ and *L-Mc1r*$^{-/-}$ mice. Values are mean ± SEM, mice n = 4-5 mice per group in each graph. **P<0.05* for the indicated comparisons by Student's t test. CA indicates cholic acid; CDCA, chenodeoxycholic acid; UDCA, ursodeoxycholic acid; MCA, muricholic acid; DCA, deoxycholic acid; LCA, litocholic acid; HDCA, hyodeoxycholic acid.

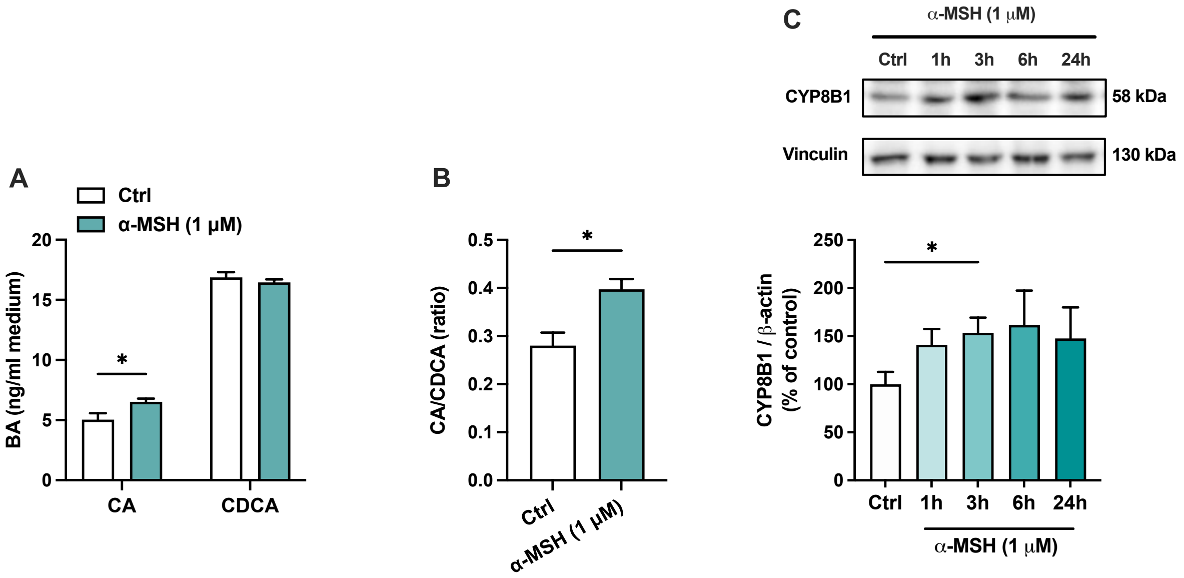


**Supplementary Figure 5.** The effects of α-MSH on bile acid production in HepG2 cells. (**A**) Quantification of cholic acid (CA) and chenodeoxycholic acid (CDCA) in the culture medium of HepG2 cells treated with 1 μM α-MSH for 24 hours. **(B)** The ratio of CA to CDCA in the culture medium of HepG2 cells treated with 1 μM α-MSH for 24 hours. n = 3-4 per group in each graph. (**C**) Representative Western blots and quantification of CYP8B1 in HepG2 cells treated with 1 μM α-MSH for 1, 3, 6 or 24 hours. n = 4-6 per group.Values are mean ± SEM, **P<0.05* for the indicated comparisons by Student's t test (**A** and **B**) or by one-way ANOVA and Dunnet *post hoc* tests (**C**).

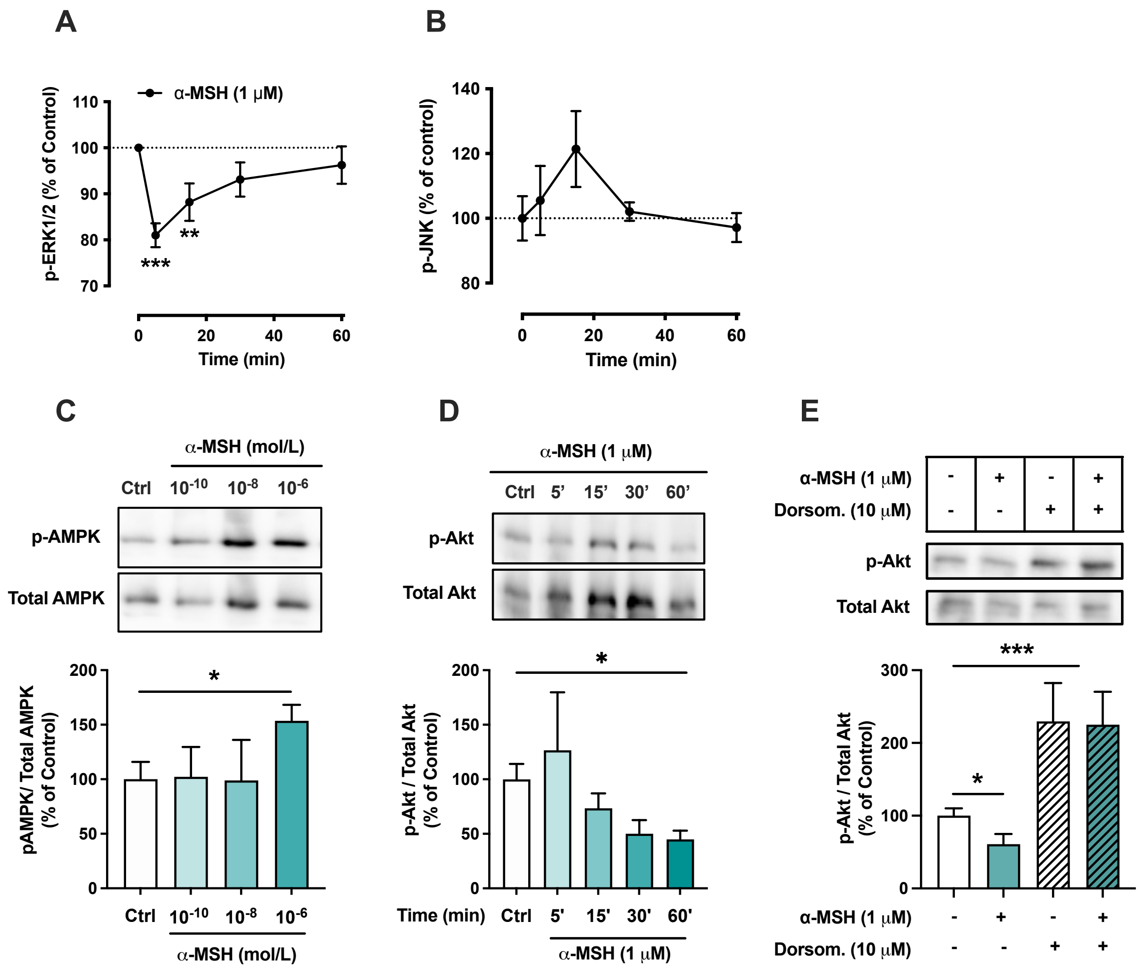


**Supplementary Figure 6.** The effects of α-MSH on the phosphorylation of ERK1/2, JNK, AMPK and Akt in HepG2 cells. **(A** and **B)** Quantification of phosphorylated ERK1/2 and JNK by ELISA assays in HepG2 cells treated with 1 µM α-MSH for 5, 15, 30 or 60 min. ***P<0.01* and ****P<0.001* versus Control (0 min). **(C)** Representative Western blots and quantification of phosphorylated AMPK level (p-AMPK normalized against total AMPK) in HepG2 cells treated with different concentrations of α-MSH (0.1 nM, 10 nM or 1 µM) for 5 minutes. (**D**) Representative Western blots and quantification of phosphorylated Akt level (p-Akt normalized against total Akt) in HepG2 cells treated with 1 µM α-MSH for 5, 15, 30 or 60 min. (**E**) Representative Western blots and quantification of phosphorylated AMPK level (p-AMPK normalized against total AMPK) in HepG2 cells treated with 1 µM α-MSH for 60 min in the presence or absence of the AMPK inhibitor dorsomorphin (1 µM). Values are mean ± SEM, **P<0.05, **P<0.01* and ****P<0.001* for the indicated comparisons by one-way ANOVA and Dunnet *post hoc* tests (**A-D**) or by two-way ANOVA and Tukey *post hoc* tests (**E**). n = 4-6 per group in each graph.

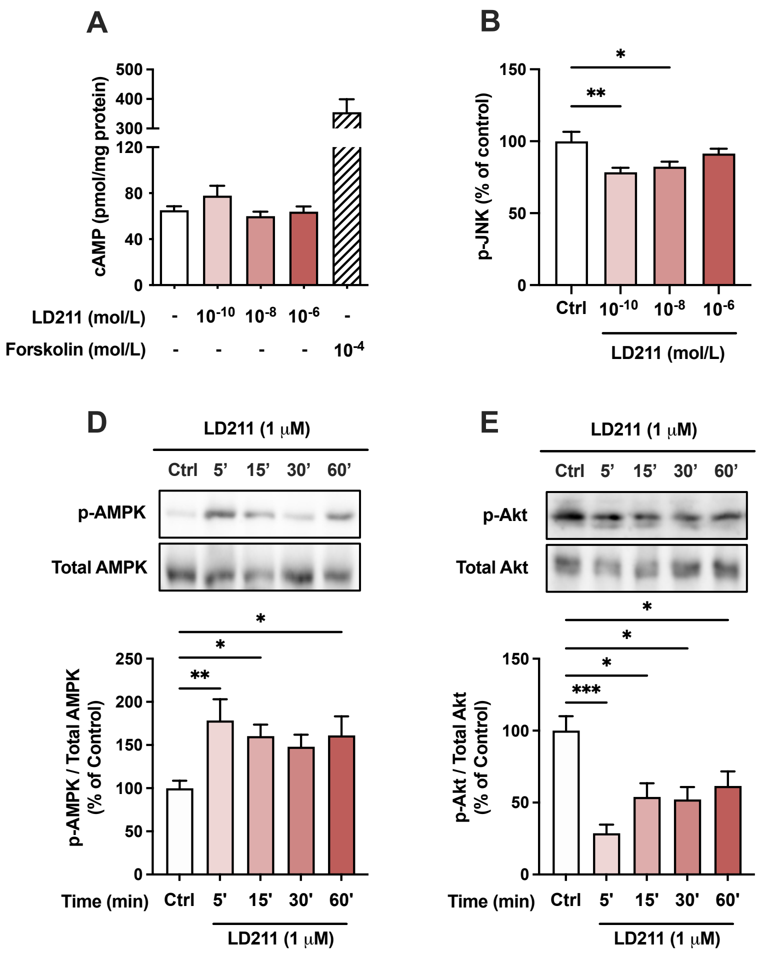


**Supplementary Figure 7.** The effects of the selective MC1-R agonist LD211 on intracellular signaling pathways in HepG2 cells. **(A)** Quantification of intracellular cAMP level in HepG2 cells treated with different concentrations of LD211 (0.1 nM, 10 nM or 1 μM) for 30 min. The adenylyl cyclase activator forskolin (10 μM) was used as a positive control. **(B** and **C)** Quantification of phosphorylated ERK1/2 and JNK by ELISA assays in HepG2 cells treated with different concentrations of LD211 (0.1 nM, 10 nM or 1 μM) for 10 min. **(D)** Representative Western blots and quantification of phosphorylated AMPK level (p-AMPK normalized against total AMPK) in HepG2 cells treated with 1 μM LD211 for 5, 15, 30 or 60 minutes. (**E**) Representative Western blots and quantification of phosphorylated Akt level (p-Akt normalized against total Akt) in HepG2 cells treated with 1 μM LD211 for 5, 15, 30 or 60 min. Values are mean ± SEM, **P<0.05, **P<0.01* and ****P<0.001* for the indicated comparisons by one-way ANOVA and Dunnet *post hoc* tests. n = 4-6 per group in each graph.
